## Supplemental Text and Figures for "Evolutionary insights into primate skeletal gene regulation using a comparative cell culture model"

All data processing and analytical scripts can be found at: https://github.com/ghousman/human-chimp-skeletal-scRNA.

### Supplemental Methods

**Validation of MSC Differentiation**
The differentiation of iPSC lines into MSCs was validated using criteria outlined by the International Society for Cell Therapy (**ISCT**) [1]. Specifically, MSCs must be plastic-adherent, express a defined set of cell surface markers, and display osteogenic, adipogenic, and chondrogenic potential. These features were assessed in human and chimpanzee iPSC-derived MSCs that were used in this study, and results confirm that iPSCs were successfully differentiated into MSCs.

*Morphological Changes*
The morphology of iPSCs and MSCs were observed by phase contrast imaging using an EVOS M5000 Imaging System (Thermo Fisher Scientific, Waltham, MA, USA). As expected, all MSC lines are plastic adherent and show characteristic elongated shape, as opposed to the more rounded shape of iPSCs (**Figure S1.1A**).

*Cell Surface Marker Changes*
Differentiation of iPSCs into MSCs was validated using flow cytometry targeting MSC markers that have been previously described by the ISCT criteria [1]. Specifically, the MSC Analysis Kit (562245, BD Biosciences, San Jose, CA, USA) was used to detect the presence or absence of CD90, CD73, CD105, CD45, CD34, CD14 or CD11b, CD19, and HLA-DR surface molecules. This kit was combined with the Zombie Violet Fixable Viability Kit (423113, BioLegend, San Diego, CA, USA) in order to also account for cell viability.

Triplicates of each iPSC and MSC line were stained according to manufacturer instructions, and flow cytometry was performed on a BD Bioscience LSRII at the Cytometry and Antibody Technology Facility (RRID: SCR_017760) at the University of Chicago. Data were analyzed using BD FACS Diva software (BD Biosciences) and FCS Express 6 (De Novo Software, Pasadena, CA, USA). As expected, iPSCs express CD90, MSCs express CD90, CD73, and CD105, and neither express CD45, CD34, CD14 or CD11b, CD19 and HLA-DR (**Figure S1.1B-C, Table S5**). This is consistent with criteria outlined by the ISCT [1].

*Osteogenic, Chondrogenic, and Adipogenic Potential*
iPSC-derived MSCs underwent osteogenic, chondrogenic, and adipogenic differentiations using protocols modified from [2] in order to assess their potential to differentiate into each of these lineages. The osteogenic differentiation protocol is described in the manuscript, and the other protocols are described as follows.

For adipogenic differentiations, iPSC-derived MSCs were cultured to at least passage 6 and seeded at 6.8 x $10^4$ cell/$cm^2$ in uncoated polystyrene culture dishes (4-10cm diameter). After culturing cells for 1 day in MSC medium, this medium was replaced with adipogenic medium that contained DMEM (11965-092, Thermo Fisher Scientific) supplemented with 10% FBS (FB5001, Thomas Scientific, Swedesboro, NJ, USA), 1% Penicillin/Streptomycin (30-002-CI, Corning, Bedford, MA, USA), 2mM GlutaMax (35050-061, Thermo Fisher Scientific), 10ug/mL Insulin (I2643, Sigma-Aldrich, St Louis, MO, USA), 500uM 3-Isobuthy-1-Methylxanthine (I5879, Sigma-Aldrich), 100uM Indomethacin (I8280, Sigma-Aldrich), and 100nM Dexamethasone (D4902 or D1756, Sigma-Aldrich). Cells were cultured at 37˚C in 5% $CO_2$ and atmospheric $O_2$, and medium was changed every 2-3 days for 3 weeks.

For chondrogenic differentiations, iPSC-derived MSCs were cultured to at least passage 6 and seeded at 5.0 x $10^4$ cell/$cm^2$ in culture dishes (4-10cm diameter) that were coated in 5ug/$cm^2$ Type I Collagen (sc-136157, Santa Cruz Biotechnology, Dallas, TX, USA). After culturing cells for 1 day in MSC medium, this medium was replaced with chondrogenic medium that contained DMEM (11965-092, Thermo Fisher Scientific) supplemented with 50ug/mL Vitamin C (A4034 or A8960, Sigma-Aldrich), 2mM GlutaMax (35050-061, Thermo Fisher Scientific), 40ug/mL L-Proline (P5607, Sigma-Aldrich), 100ug/mL Sodium Pyruvate (11360-070, Thermo Fisher Scientific), 100nM Dexamethasone (D4902 or D1756, Sigma-Aldrich), 1X ITS Premix (354352, Corning, Bedford, MA, USA), 10ng/mL TGF-β3 (243-B3, R&D Systems, Minneapolis, MN, USA). Cells were cultured at 37˚C in 5% $CO_2$ and atmospheric $O_2$, and medium was changed every 2-3 days for 3 weeks.

Validation of osteogenic, chondrogenic, and adipogenic differentiation potential is described in the following sections.

**Validation of Multipotent Cell Differentiation Potential using Histology**
MSCs, osteogenic cells, adipogenic cells, and chondrogenic cells were fixed using 4% paraformaldehyde in phosphate-buffered saline (sc-281692, Santa Cruz Biotechnology) and stored at 4°C until ready for staining. Once stained, cells were imaged using an Olympus SZX7 Stereo Microscope (Olympus, Center Valley, PA, USA) or an EVOS M5000 Imaging System (Thermo Fisher Scientific).

*Validation of Osteogenic Differentiation using Alizarin Red Staining*
Fixed osteogenic cells and MSCs were washed with 1X PBS and stained with Alizarin Red S solution (20mg/mL in DI water, pH 4.1-4.3; A5533, Sigma-Aldrich) for 45 minutes at room temperature. The stain was removed, and cells were washed with DI water to remove the excess stains prior to imaging. Alizarin Red S stains calcium deposits a red-orange color (**Figure 1D, Figure S1.1D, Figure S1.2A**). After imaging, stain was extracted by adding a hexadecylpyridinium chloride solution and agitating for 1 hour on a shaker. The absorbance (562nm; 655090, Greiner Bio-One, Monroe, NC, USA) was measured for replicates of the extracted stain in order to quantify variation in calcium production (**Table S6**). Statistical significance between osteogenic and MSC absorbance levels was calculated within each species (**Figure S1.1D**) and individual (**Figure S1.2B**) using one-sided Mann Whitney tests (see 10x-view-validation-data.Rmd for code). In general, calcium production is greater in osteogenic cells than in MSCs across species and individuals.

*Validation of Adipogenic Differentiation using Oil Red Staining*
Fixed adipogenic cells and MSCs were washed with 1X PBS, dehydrated briefly with ethanol, and stained with Oil Red O (5mg/mL in isopropanol; 01794 Chem-Impex International Inc., Bensenville, IL, USA) for 15 minutes at room temperature. The stain was removed, and cells were briefly dehydrated with ethanol before being washed with DI water to remove the excess stains prior to imaging. Oil Red O stains intracellular lipids and fat vacuoles a red color (**Figure S1.1D**). After imaging, stain was extracted by adding isopropanol, incubating 5 minutes at room temperature, and vortexing 15-30 seconds. The absorbance (510nm; 655090, Greiner Bio-One) was measured for replicates of the extracted stain in order to quantify variation in lipid production (**Table S7**). Statistical significance between adipogenic and MSC absorbance levels was calculated within each species (**Figure S1.1D**) using one-sided Mann Whitney tests (see 10x-view-validation-data.Rmd for code). In general, lipid production is greater in adipogenic cells than in MSCs across species and individuals.

*Validation of Chondrogenic Differentiation using Alcian Blue Staining*
Fixed chondrogenic cells and MSCs were dehydrated using a series of ethanol washes and then rehydrated with DI water before staining with Alcian Blue (5mg/mL in 0.1M HCl; A3157, Sigma-Aldrich) for 30 minutes at room temperature. The stain was removed, and cells were washed with DI water to remove the excess stains prior to imaging. Alcian Blue stains proteoglycans, which are found in connective tissue like cartilage, a blue color. In general, proteoglycan production is greater in chondrogenic cells than in MSCs (**Figure S1.1D**).

**Validation of Cell Differentiation Potential using qPCR**
Quantitative real-time PCR (**qPCR**) was also used to confirm the osteogenic, adipogenic, and chondrogenic differentiation potential of iPSC-derived MSCs. Gene expression levels of MSCs and differentiated cells were assessed for several cell type markers (**Table S8**). Primer sequences are provided in the following table.

| Gene | Forward Primer Sequence (5' to 3') | Reverse Primer Sequence (5' to 3') |
|---|---|---|
| COL1A1 | AGGGCCAAGACGAAGACATC | AGATCACGTCATCGCACAAC |
| BGLAP | CGCCTGGGTCTCTTCACTAC | CTCACACTCCTCGCCCTATT |
| RUNX2 | CAGTAGATGGACCTCGGGAA | CCTAAATCACTGAGGCGGTC |
| FABP4 | ACTGGGCCAGGAATTTGACG | CCCCATCTAAGGTTATGGTGCTC |
| PPARG | TCAGGTTTGGGCGGATGC | TCAGCGGGAAGGACTTTATGTATG |
| COL2A1 | GGCAATAGCAGGTTCACGTACA | CGATAACAGTCTTGCCCCACTT |
| SOX9 | GTACCCGCACTTGCACAAC | TCTCGCTCTCGTTCAGAAGTC |
| COL11A1 | TGGTGATCAGAATCAGAAGTTCG | AGGAGAGTTGAGAATTGGGAATC |
| GAPDH | ATGGGGAAGGTGAAGGTCGGAGTCA | TCCTGGAAGATGGTGATGGGATTTC |
| GUSB | CTCATTTGGAATTTTGCCGATT | CCGAGTGAAGATCCCCTTTTTA |

Briefly, total cellular RNA was extracted from samples using the ZR-Duet™ DNA/RNA MiniPrep (D7001, Zymo Research, Irvine, CA, USA) with a DNase (E1010, Zymo Research) treatment, and cDNA was prepared using the Maxima FS cDNA kit for RT-PCR (K1642, Thermo Fisher Scientific). qPCRs were run using the PowerUp™ SYBR™ Green Master Mix (A25742, Thermo Fisher Scientific) on an Applied Biosystems QuantStudio 6 Flex Real-Time PCR System. Expression data were collected as Ct values and relative quantification levels were calculated using the $E^{-\Delta\Delta C_T}$ method [3] which determines the deviation between sample replicate Ct values and a control sample Ct value (MSCs), normalizes the target gene to a reference gene (average expression of GAPDH and GUSB), and accounts for the qPCR efficiency of each gene calculated using in-house serial dilution standards. Statistical significance between the relative quantification levels osteogenic, adipogenic, and chondrogenic cells and those of MSCs was calculated across individuals and species (**Figure S1.1E**) using one-sided t-tests on ΔΔCt values (see 10x-view-validation-data.Rmd for code).

**Cell Dissociations for Single-Cell RNA Sequencing**

Cells used for single-cell RNA sequencing were first washed with 1X PBS (21-040-CV, Thermo Fisher Scientific) and then dissociated from adherent conditions into single-cell suspensions. For iPSCs, cells were incubated in a dissociation reagent (0.25mM EDTA, 150mM NaCl in PBS) for 2-5 minutes, pelleted at 1000 rpm for 5 minutes, and resuspended in a buffer solution (561527, BD Biosciences). For MSCs, cells were incubated in 0.05% Trypsin (25-052-CI, Corning) for 5-7 minutes. Once the cells detached, trypsin was neutralized with MSC medium, and the cells were pelleted at 1000 rpm for 5 minutes and resuspended in a buffer solution (561527, BD Biosciences). For osteogenic cells, cells were first incubated in 1mg/mL collagenase II (LS004204, Worthington Biochemical Corporation, Lakewood, NJ, USA) in 1X HBSS (21-022-CV, Thermo Fisher Scientific) for 10 minutes. Collagenase II was neutralized with MSC medium and removed. After a second 1X PBS wash, cells were incubated in 0.25% Trypsin (25-053-CI, Corning) for 10 minutes. Once the cells detached, trypsin was neutralized with MSC medium, and the cells were pelleted at 1000 rpm for 5 minutes and resuspended in a buffer solution (561527, BD Biosciences).

**Assigning 10X Single-Cell Barcodes to Species**

Single-cell 10X cell barcodes were bioinformatically reassigned to their species of origin using standard 10X Genomics Cell Ranger 3.1.0 pipelines (Cell Ranger) [4], with the exception that reads were aligned to both the human genome (hg38) and the chimpanzee genome (panTro6) and a curated set of orthologous exons [5] was used for transcriptome alignment. Briefly, Cell Ranger aligns reads to each of two genomes, retaining only those reads that specifically align to one genome and discarding reads that align to both genomes. Each cell barcode is then classified as human or chimpanzee based on which genome has more aligned UMI counts or as a multiplet if the human UMI counts and the chimpanzee UMI counts exceed the $10^{th}$ percentile of each species' distribution.

Because Cell Ranger was optimized for mixtures of human and mouse cells, which are more genetically divergent than humans and chimpanzees, we performed additional tests to determine how accurate Cell Ranger is at assigning human and chimpanzee cells and identified more accurate cell assignment criteria that we used for our data. Specifically, we ran Cell Ranger as described above on one dataset from our group that contained only human cells [YG-AH-2S-ANT-1] [6], one dataset that contained only chimpanzee cells from an external group [SRR8403265] [7], and one dataset that contained only human cells from the same external group [SRR8403264] [7]. Cell Ranger assigns most of the cell barcodes in these datasets as multiplets (**Figure S2.2A-C**) even though the majority of UMI counts resulting from the human-only datasets were specific to the human genome and the majority of UMI counts resulting from the chimpanzee-only dataset were specific to the chimpanzee genome (**Figure S2.2D-F**). On average, about 3% of UMIs from the human-only dataset align to the chimpanzee genome and about 4% of UMIs from the chimpanzee-only dataset align to the human genome (**Figure S2.2D-F**). We were able to correctly assign all cells in these datasets when considering the ratio of human-aligned UMIs to the total number of aligned UMIs within each cell. Specifically, using a ratio cutoff of 0.94 successfully assigned all cells from the human-only datasets as human, and using a ratio cutoff of 0.09 successfully assigned all cells from the chimpanzee-only datasets as chimpanzees (**Figure S2.2G-I**). These cutoffs account for the chance that human reads may align to the chimpanzee genome and that chimpanzee reads may align to the human genome due to high sequence similarity and differences in reference annotation qualities. Thus, they are the minimum criteria for assigning human and chimpanzee cells in single-cell mixtures.

However, for actual human and chimpanzee single-cell mixtures, it is also expected that there will be additional background human and chimpanzee reads due to cells that lyse before droplet formation. This background will vary from batch to batch, making it difficult to explicitly calculate. Thus, for the data collected in this study, we opted to slightly increase the ratio cutoffs to 0.9 and 0.1 for humans and chimpanzees, respectively. Using these cutoffs greatly improves species assignments as compared to those produced directly from Cell Ranger (**Figure S2.3**) (see 10x-data-0-process-test.Rmd for code).

**Single-Cell Data Processing**

As described above, the standard Cell Ranger pipeline using both human and chimpanzee genomes to determine species assignments discards reads that align to both genomes. Because the human and chimpanzee genomes have a high sequence similarity, using this pipeline discards an extremely high number of reads, making downstream data analyses extremely difficult.

Thus, to improve the number of reads retained for downstream analyses, we performed two additional runs of the Cell Ranger pipeline such that raw single-cell RNA sequencing reads were either aligned to the human genome (hg38) or the chimpanzee genome (panTro6) and then subsequently aligned to a curated set of orthologous exons [5] (**Figure S2.1D**) (see 10x-data-1-proces.Rmd for code). Specifically, for 10X barcodes that were assigned human as described above, reads were aligned to the human genome and orthologous exons to produce gene count matrices. Alternatively, for 10X barcodes that were assigned chimpanzee as described above, reads were aligned to the chimpanzee genome and orthologous exons to produce gene count matrices.

These matrices containing gene counts per 10X barcode were then combined and further processed using the Seurat package (v3.1.2) [8,9]. In particular, cells assigned as multiplets were removed, and cells were filtered if fewer than 1000 UMIs were detected, if fewer than 700 genes were detected, and if more than 25% of reads mapped to the mitochondria. This resulted in a total of 101,000 cells that were used in subsequent analyses (see 10x-data-1-process.Rmd and 10x-data-2-qc.Rmd for code).

**Single-Cell Data Integration**

After processing (see 10x-data-1-process.Rmd for code) and filtering (see 10x-data-2-qc.Rmd for code), data were integrated using Seurat (v3.1.2) [8,9] (see 10x-data-3-integrate.Rmd for code). This involved a series of steps, starting with data normalization. Data were log-normalized using the NormalizeData function in Seurat, which divides gene counts by the total counts, multiplies them by a scale factor of 10,000, and then natural-log transforms them.

Following this, cell cycle scores were calculated using the CellCycleScoring function in Seurat. Specifically, cell cycle scores were assigned to cells based on G2/M and S phase markers in each cell. The calculations depend on the average gene expression levels across all the cells in a dataset, so data subsetting was an important consideration. Because we anticipated that cell cycle would vary across different time points (iPSCs and MSCs continuing to replicate while late stage osteogenic cells would halt replication), cell cycle scores were calculated separately in datasets comprising cells grouped by cell line and cell type (n=42).

Finally, in order to classify cell types across collections and species, data were integrated. Integration was performed because simple data merging retained unwanted technical variation and did not fully integrate species. Even merging datasets using the intersect of the most variable genes in humans and chimpanzees did not fully integrate data across species. Thus, data were instead integrated in Seurat.

Successful data integration is dependent on how the data are subsetted and integrated across. In order to remove unwanted technical variation and allow for the successful calling of cell types using biological variation of interest, we decided to use the datasets described above (n=42) but merge those coming from the same cell line and replicate (n=14). Thus, an individual dataset comprised all cells (iPSCs, MSCs, and osteogenic cells) derived from an individual cell line replicate. Additional subsetting schemes were considered, including grouping cells by species (n=2), grouping cells by species of origin and cell type (n=6), grouping cells by 10X collection batch (n=7), grouping cells by 10X collection batch and cell type (n=21), and grouping cells by cell line and cell type (n=42). However, these alternative schemes result in cell integrations that do not fully remove batch and individual effects.

Specifically, datasets were integrated across individuals and collection batches using the reference-based, reciprocal PCA method in Seurat. This method, while perhaps less accurate than canonical correlation analysis, is computationally efficient, which was necessary for the large number of cells considered in this study. In this method, reciprocal PCA is performed to identify an effective space in which to find anchors. Specifically, each dataset is projected into every other dataset's PCA space, and anchors are constrained by the same mutual neighborhood requirement. We defined integration anchors as all genes that had non-zero UMI counts (n=18,482), as this resulted in better integration than other gene subsets tested. Because subsets of databases are assigned as references, each dataset is compared to the references instead of performing all pairwise comparisons. For these data, H1-r2 and C1-r2 were defined as references because they differentiated well based on standard validation metrics and they served as replicate samples in the study. During integration, the anchors between datasets (which represent pairwise correspondences between individual cells) were then used to harmonize the datasets (transfer information from one dataset to another).

Although a substantial amount of unwanted variation was removed from these integrated data, integrated data reductions were still correlated with UMI counts and percent of mitochondrial reads. Thus, this variation was regressed out using the ScaleData function in Seurat. Cell cycle phase was also correlated with integrated data reductions, but this variation was not regressed out because this is not recommended for differentiating cells.

**Single-Cell Data Annotation**

Cell type classifications were defined and assigned to single-cell data using several methods (see 10x-data-4-classification-tot.Rmd and 10x-data-plot-classification.Rmd for code). The methods reported in the manuscript include the stage of differentiation at which cells were collected, unsupervised clustering, and ad hoc assignments. Additionally, modifications to parameters in each of these classification schemes were also examined, as well as several other classification schemes.

*Unsupervised Clustering*

For unsupervised clustering (see 10x-data-plot-classification-clusters.Rmd for code) beyond the resolutions of 0.05, 0.25, and 0.50 that are discussed in the manuscript, resolutions of 0.10, 0.75, and 1.00 were also considered. Additionally, these higher resolutions were used to isolated osteogenic-specific clusters. In particular, a resolution of 0.50 identifies 4 osteogenic clusters, and a resolution of 0.75 identifies 8 osteogenic clusters. Both of these annotations were considered in downstream analyses.

*Ad Hoc Assignments*

For ad hoc assignments (see 10x-data-plot-classification-adhoc.Rmd for code), candidate gene expression patterns were used to determine general ad hoc assignments (iPSC, MSC, or osteogenic) and osteogenic ad hoc assignments (preosteoblasts, osteoblasts, embedding osteoblasts, mineralizing osteoblasts, maturing osteocytes). In both instances, a cell was defined as positively expressing a particular gene if the expression level for that gene within the cell was greater than the mean expression level of that gene across all cells. Expression levels were based on scaled integrated data that had confounding variables regressed out, as described above. Each ad hoc assignment was defined by the expression patterns of multiple genes, as described below, and for a cell to be classified to a particular ad hoc assignment, all criteria for gene expression patterns had to be met. To be thorough, additional cutoff thresholds were considered (general ad hoc assignments: mean expression level +/- 0.01, 0.05, and 0.10; osteogenic ad hoc assignments: mean expression level +/- 0.01) with more lenient thresholds resulting in more cells classified to a particular ad hoc assignment and with more conservative thresholds resulting in fewer cells classified to given ad hoc assignments.

For general ad hoc assignments, pluripotent candidate genes included *POU5F1*, *TDGF*, *SOX2*, and *EPCAM*, mesenchymal candidate genes included *THY1*, *NT5E*, *ENG*, and *CD44*, and the primary osteogenic candidate gene included *ALPL*. Pluripotent cells were defined as those expressing any of the pluripotency candidate genes (*POU5F1*, *TDGF*, *SOX2*, *EPCAM*) but no mesenchymal-specific candidate genes (*NT5E*, *ENG*, *CD44*). Mesenchymal cells were defined as those expressing any of the mesenchymal candidate genes (*THY1*, *NT5E*, *ENG*, *CD44*) but no pluripotent candidate genes

(*POU5F1*, *TDGF*, *SOX2*, *EPCAM*) or osteogenic candidate genes (*ALPL*). Finally, osteogenic cells were defined as those expressing osteogenic candidate genes (*ALPL*) but no pluripotent candidate genes (*POU5F1*, *TDGF*, *SOX2*, *EPCAM*). Additional osteogenic candidate genes (*COL1A1* and *COL1A2*) were considered but not included because these genes are also expressed in mesenchymal cells which complicated disentangling these cells.

For osteogenic ad hoc assignments, cells had to first be assigned as osteogenic using the general ad hoc criteria described above. Then cells were classified into different stages of osteogenesis (preosteoblasts, osteoblasts, embedding osteoblasts, mineralizing osteoblasts, and maturing osteocytes) using candidate genes that are known to vary in expression levels across these stages of differentiation [10]. These genes include *RUNX2*, *BGLAP*, *CAPG*, *DMP1*, *PHEX*, *MEPE*, *E11*, *SOST*, and *HYOU1*. *E11* is not a part of the curated orthologous exon set used in this study, so it was not considered for osteogenic ad hoc assignments. Additionally, some of these genes did not seem appropriate for determining classifications. Thus, we examined several different combinations of genes ranging from a combination of 8 genes which was most consistent with literature [10] to an intermediary combination of 6 genes, and lastly to a combination of 4 genes which was the simplest definition considered.

First, we examined assignments using all 8 genes that overlap with our data (**Figure S4.3A**). Preosteoblasts were defined as cells expressing *RUNX2* but no other osteogenic genes. Osteoblasts were defined as cells expressing *BGLAP*, having variable expression of *RUNX2*, and not expressing other osteogenic genes. Embedding osteoblasts were defined as cells expressing either *CAPG*, *DMP1*, or *PHEX*, having variable expression of *RUNX2* and *BGLAP*, and not expressing other osteogenic genes. Mineralizing osteoblasts were defined as cells expressing *MEPE*, having variable expression of *RUNX2*, *BGLAP*, *CAPG*, *DMP1*, and *PHEX*, and not expressing other osteogenic genes. Maturing osteocytes were defined as cells expressing either *SOST* or *HYOU1* and having variable expression of all other osteogenic genes. All other cells were defined as unknown osteogenic.

Next, we examined assignments using only 6 of these genes (**Figure S4.3C**). In this set, *CAPG* was removed because it is highly expressed all cells in our data (pluripotent, mesenchymal, and osteogenic cells), which diminishes its usefulness as a marker for differentiating subtle cell types. Additionally, *HYOU1* was removed because it is a heat-shock protein. While this is a potentially useful marker for maturing osteocytes *in vivo* which are within a hypoxic environment of mineralized bone matrix, its expression within *in vitro* osteogenic cells may be affected by the oxygen and temperature changes that are inherent to cell culture environments. Using this combination of 6 genes, preosteoblasts were again defined as cells expressing *RUNX2* but no other osteogenic genes. Osteoblasts were again defined as cells expressing *BGLAP*, having variable expression of *RUNX2*, and not expressing other osteogenic genes. Embedding osteoblasts were defined as cells expressing either *DMP1* or *PHEX*, having variable expression of *RUNX2* and *BGLAP*, and not expressing other osteogenic genes. Mineralizing osteoblasts were defined as cells expressing *MEPE*, having variable expression of *RUNX2*, *BGLAP*, *DMP1*, and *PHEX*, and not expressing other osteogenic genes. Maturing osteocytes were defined as cells expressing *SOST* and having variable expression of all other osteogenic genes. All other cells were defined as unknown osteogenic.

Lastly, we examined assignments using only 4 of these genes (**Figure S4.3E**). In this set, *DMP1* and *SOST* were removed because of their substantially lower expression levels in osteogenic cells as compared to other candidate genes, and in the case of *DMP1*, its redundant nature with *PHEX*. Using this combination of 4 genes, preosteoblasts were again defined as cells expressing *RUNX2* but no other osteogenic genes. Osteoblasts were again defined as cells expressing *BGLAP*, having variable expression of *RUNX2*, and not expressing other osteogenic genes. Embedding osteoblasts were defined as cells expressing *PHEX*, having variable expression of *RUNX2* and *BGLAP*, and not expressing other osteogenic genes. Mineralizing osteoblasts were defined as cells expressing *MEPE* and having variable expression of *RUNX2*, *BGLAP*, and *PHEX*. Maturing osteocytes were defined as cells expressing *MEPE*, not expressing *RUNX2* and *BGLAP*, and having variable expression of all other osteogenic genes. All other cells were defined as unknown osteogenic.

Because transitioning between stages of osteogenesis is gradual, we felt that flexible cell type definitions which allow for variable expression patterns of stage-specific genes that are superseded in subsequent osteogenesis stages would be more biologically relevant for interpreting a differentiation trajectory and would be better able to classify cell types. Nevertheless, for each combination of genes, we also considered more strict definitions of each cell type that did not allow for variable gene expression. These definitions reduced the overall number of cell classifications as shown in the table below and were not considered further.

| Gene Set Combination | Preosteoblasts | Osteoblasts | Embedding Osteoblasts | Mineralizing Osteoblasts | Maturing Osteocytes | Unknown Osteogenic |
|---|---|---|---|---|---|---|
| 4 genes (flexible) | 2004 | 3878 | 3019 | 4473 | 2443 | 1758 |
| 6 genes (flexible) | 1509 | 1664 | 6477 | 6733 | 193 | 999 |
| 8 genes (flexible) | 273 | 372 | 6016 | 4682 | 6030 | 202 |
| 4 genes (strict) | 2004 | 1909 | 969 | 86 | 313 | 1758 |
| 6 genes (strict) | 1509 | 1020 | 755 | 45 | 45 | 999 |
| 8 genes (strict) | 273 | 215 | 362 | 25 | 0 | 202 |

As shown in the above table, for our flexible cell type definitions, using a combination of 8 genes resulted in more cells assigned to late stages of osteogenesis than the other gene combination sets, a combination of 6 genes resulted in more cells assigned to intermediate stages of osteogenesis than the other gene combination sets, and a combination of 4 genes resulted in more cells assigned to early stages of osteogenesis than the other gene combination sets. Our expectation was that most osteogenic cells would be in the early stages of osteogenesis, and the osteogenic ad hoc assignments determined using a combination of 4 genes most closely matched this expectation.

To formally assess which combination of osteogenic genes determined the most biologically meaningful classifications, we examined the pairwise correlations between the whole transcriptome pseudobulk of each osteogenic ad hoc classification with the assumption that correlations should be greatest between adjacent stages of osteogenesis and decrease as the separation between stages increase (e.g., correlation should be greatest between preosteoblasts and osteoblasts, reduced between preosteoblasts and embedding osteoblasts, further reduced between preosteoblasts and mineralizing osteoblasts, and the lowest between preosteoblasts and maturing osteocytes). This pattern was only observed for osteogenic ad hoc assignments determined using 4 genes (*RUNX2*, *BGLAP*, *PHEX*, *MEPE*) (**Figure S4.3B,D,F**) Of note, all the pairwise correlations are generally high, because these cell types are not substantially separated across cell differentiation space. This is in contrast to pairwise correlations between the whole transcriptome pseudobulk of each stage of differentiation (**Figure S2.5A**), unsupervised clusters at a resolution of 0.05 (**Figure S2.5B**), or general ad hoc assignments (**Figure S2.5C**). Based on these additional analyses, the final set of candidate genes used for defining osteogenic ad hoc assignments included *RUNX2*, *BGLAP*, *PHEX*, and *MEPE*, and these data are discussed further in the manuscript.

**Topic Modeling of Single-Cell Data**

Topic modeling (see 10x-data-4-classification-tot.Rmd, 10x-data-plot-classification-gom.Rmd, and 10x-data-plot-classification.Rmd for code) was performed as an alternative method to examined data structure. These methods are described in the manuscript.

**Differential Expression Analyses**

Differential expression (DE) was performed using standard tests in the dream package [11] (see 10x-data-5-diffexp.Rmd and 10x-plot-diffexp.Rmd for code) and a Bayesian clustering approach implemented by Cormotif [12] (see 10x-data-5-cormotif-tot-cnts_v2.Rmd and 10x-plot-cormotif-tot-cnts_v2.Rmd for code). These methods are described in the manuscript. Briefly, for both analyses, pseudobulk expression values were calculated as described in the manuscript, and mitochondrial genes, ribosomal genes, and genes that had an average log2 transformed CPM less than or equal to zero were removed from the data (**Table S10**). Additionally, for standard DE tests, we also randomly subsampled cell classifications to equal numbers across species before calculating pseudobulk gene expression values and performing DE tests (see 10x-data-5-diffexp.Rmd for code). This was done to ensure that the number of cells assigned to each classification did not affect broad trends that we detected in our DE results.

Genes were classified as significantly DE if they had a false discovery rate (**FDR**) < 0.01 for standard tests or if they had a posterior probability > 0.65 for Cormotif tests. These thresholds were selected after testing several different cutoffs (FDR < 0.50, 0.40, 0.30, 0.20, 0.10, 0.05, 0.01, or 0.001; posterior probability > 0.50, 0.55, 0.60, 0.65, 0.70, 0.75, 0.80,

0.85, 0.90, 0.925, 0.95, 0.975, or 0.99) to call interspecific DE genes within stages of differentiation (see 10x-plot-diffexp-thresh.Rmd and 10x-plot-cormotif-cnts-thresh_v2.Rmd for code). For each set of interspecific DE genes identified, we performed gene set enrichment tests as described below using relevant external gene sets (**Table S11**). We expected that previously identified interspecific DE genes in iPSCs [13,14] should only be enriched among genes we identified as interspecific DE in pluripotent cells (Time 0) and that previously identified interspecific DE genes in alternative cell and tissue types (non-pluripotent, non-mesenchymal, and non-osteogenic) [13,15] should not be enriched among any interspecific DE genes identified in this study. The FDR and posterior probability thresholds that best matched these enrichment expectations (**Figure S3.3, Figure S3.6**) were then used for downstream analyses (FDR < 0.01; posterior probability > 0.65).

**Gene Expression Variance**

We examined whether variance in gene expression differs across cell classifications (see 10x-data-6-variance-tot.Rmd for code) as described in the manuscript.

**Concordance and Functional Enrichment Tests**

We assessed the enrichment of Gene Ontology categories among various sets of genes identified in this study using the enrichGO function in the clusterProfiler v3.12.0 package in R [16] (see 10x-plot-classification.Rmd and 10x-plot-enrichment-gom.Rmd for code) and the enrichment of relevant gene sets (**Table S11**) in the DE genes in this study using a Fisher's exact test (see 10x-plot-diffexp.Rmd and 10x-plot-cormotif- cnts_v2.Rmd for code) as described in the manuscript. Because very few skeletal-trait-associated genes are known for hip geometry [17] and bone area [18], we also examined which of these skeletal-phenotype genes overlap our interspecific DE genes. In particular, out of the 10 genes associated with GWAS hits for hip geometry [17], only seven passed filtering criteria and were tested in our data (*FGFR4, NSD1, LRP5, PPP6R3, GAL, CCDC91, RUNX1*), with *FGFR4* only meeting these criteria for Cormotif DE tests within stages of differentiation. Additionally, out of the 15 genes associated with GWAS hits for bone area [18], only eight passed filtering criteria and were tested in our data (*BCKDHB, COL11A1, CTDSP2, DYM, HHIP, SH3GL3, SOX9, WNT4*), with only *BCKDHB, COL11A1, CTDSP2,* and *DYM* meeting these criteria for all of DE tests. Specifically, *HHIP* was not examined in any standard DE tests, *SH3GL3* was not examined in any standard DE tests or Cormotif DE tests within stages of osteogenesis, *SOX9* was not examined in standard DE tests within stages of osteogenesis, and *WNT4* was not examined in standard or Cormotif DE tests within stages of osteogenesis.

#### Supplemental Results

##### Confirming multipotent differentiation potential of MSCs

Osteogenic, adipogenic, and chondrogenic differentiations were performed to confirm the multipotent differentiation potential of MSCs. Validation of osteogenic differentiation via Alizarin Red staining of calcium deposits (**Figure 1D, Figure S1.1D, Figure S1.2**) is discussed in the manuscript. Additionally, adipogenic differentiation was validated using Oil Red staining which confirmed the production of fat globules, and chondrogenic differentiation was validated using Alcian Blue staining which confirmed the production of collagen extracellular matrix (**Figure S1.1D**). As a secondary confirmation, qPCR was used to confirm the increased expression of *COL1A1, BGLAP*, and *RUNX2* in osteogenic cells, *FABP4* and *PPARG* in adipogenic cells, and *COL2A1*, *SOX9*, and *COL11A1* in chondrogenic cells (**Figure S1.1E, Table S8**).

##### Evaluating single-cell data balance and reproducibility

Species are fairly well integrated within each stage of differentiation (**Figure S2.4A**). There is some separation, but other known batch effects (**Table S2, Table S9**) do not seem to account for these interspecific cell distribution separations. Using the gene expression patterns recovered from different technical replicates, we also found that these scRNA-seq data are highly reproducible. Specifically, we observed strong correlations between replicates when considering gene expression within each stage of differentiation, both when species are combined (**Figure 2E-G**) or analyzed separately (**Figure S2.4B**). For these correlation tests, log-normalized scRNA-seq gene counts as described above were first averaged across cells in each technical replicate and then log-transformed. Lastly, Pearson correlations were calculated between the log-transformed average gene counts of one replicate and the log-transformed average gene counts of the second replicate for a given cell classification.

##### Examining alternative cell annotations as compared to stages of differentiation

We examined whether our processed and integrated scRNA-seq data remains balanced across species and reproducible, displays divergent transcriptomic patterns, and exhibits expected marker gene expression levels when alternative classification schemes, including unsupervised clustering and ad hoc assignment methods, are used to annotate cells.

Unsupervised clustering with a resolution of 0.05 identified 5 clusters – 3 pluripotent clusters (iPSC.c1, iPSC.c2, iPSC.c3), 1 mesenchymal cluster (MSC.c1), and 2 osteogenic clusters (Osteogenic.c1, Osteogenic.c2) (**Figure S2.6A**). Within each cluster, similar numbers of cells were recovered in each species (**Figure S2.6B**), and UMI counts per cell (**Figure S2.6C**) and gene counts per cell (**Figure S2.6D**) are also similar across species. Additionally, gene expression patterns recovered from different technical replicates are strongly correlated within each cluster (**Figure S2.6E**). Whole transcriptome pseudobulk correlations also show clear transcriptomic differences across cell classification (**Figure S2.5B**). Finally, GO enrichments in cluster-specific marker genes reveal expected and biologically relevant functional enrichments (**Figure S2.6F-H**).

General ad hoc assignments classified cells into iPSCs, MSCs, and osteogenic cells as described above (**Figure S2.7A**). Within each cell classification, similar numbers of cells were recovered in each species (**Figure S2.7B**), and UMI counts per cell (**Figure S2.7C**) and gene counts per cell (**Figure S2.7D**) are also similar across species. Additionally, gene expression patterns recovered from different technical replicates are strongly correlated within each cell assignment (**Figure S2.7E**). Whole transcriptome pseudobulk correlations also show clear transcriptomic differences across cell classification (**Figure S2.5C**). Finally, GO enrichments in assignment-specific marker genes reveal expected and biologically relevant functional enrichments (**Figure S2.7F-H**).

Cell groupings produced using unsupervised clustering and ad hoc assignments have characteristics very similar to those of cell groupings defined by the stage of differentiation at which cells were collected as described in the manuscript. Thus, since stage of differentiation serve as good proxies for general cell types, this classification scheme is used for all downstream analyses described in the manuscript.

**Testing for concordance and functional enrichment of interspecific DE genes across stages of differentiation**
In order to ascertain the biological functions of interspecific DE genes identified across stages of differentiation (both via standard and Cormotif analyses), we tested for the enrichment of relevant external gene datasets (**Figure 3E-F, Figure S5.1A**) and GO categories (**Figure S5.3A, Figure S5.4A, Table S12, Table S13**). Resulting trends for standard DE tests are supplied here, while details for the Cormotif DE tests are discussed in the manuscript.

As expected, previously identified interspecific DE genes in iPSCs [13,14] are enriched among genes we identified as interspecific DE in pluripotent cells (Time 0) but not among genes classified as interspecific DE in later stages of differentiation (Time 1 and Time 2). Additionally, previously identified interspecific DE genes in alternative cell types and tissues (non-pluripotent, non-mesenchymal, and non-osteogenic) [13,15] are not enriched among genes we identified as interspecific DE in this study. Two exceptions are that previously identified interspecific DE genes in kidney tissue [15] are enriched among genes we identified as interspecific DE in pluripotent cells (Time 0) and that previously identified interspecific DE genes in liver tissue [15] are enriched among genes we identified as interspecific DE across earlier stages of differentiation (Time 0 and Time 1). Lastly, previously identified interspecific DE genes in CNCCs [14] are enriched among genes we identified as interspecific DE across later stages of differentiation (Time 1 and Time 2) and among genes we identified as interspecific DE in osteogenic cells (Time 2), and differentially methylated regions (**DMRs**) in bone tissues [19] are enriched among genes we identified as interspecific DE in mesenchymal cells (Time 1).

We did not find enrichment of interspecific DE genes among relevant GO categories. Similarly, we found no overlap of skeletal trait and disease-related genes among interspecific DE genes. Specifically, we found no significant enrichment of hip geometry [17], bone area [18], height [20], osteoporosis [21], or osteoarthritis [22] related loci among interspecific DE genes identified in this study. Because there are very few genes associated with GWAS hits for hip geometry [17] and bone area [18] that also overlap with genes tested in this study, we also examined overlapping genes individually. Of the hip geometry genes [17] that were tested in our data, we found that *PPP6R3* is also a interspecific DE gene that are shared between pluripotent cells (Time 0) and mesenchymal cells (Time 1). Additionally, of the bone area genes [18] that were tested in our data, *BCKDHB* and *COL11A1* are interspecific DE genes unique to pluripotent cells (Time 0).

**Identifying DE across an alternative general cell classification system (unsupervised clustering)**
Within each general unsupervised cluster, we discovered hundreds of DE genes between species. Specifically, standard DE analyses of pseudobulk data detected 2,155 interspecific DE genes in iPSC.c1, 413 in iPSC.c2, 0 in iPSC.c3, 936 in MSC.c1, 534 in Osteogenic.c1, and 209 in Osteogenic.c2 with an FDR $< 0.01$ (**Figure S3.1A**). Focusing in on the primary cluster for each cell type (iPSC.c1, MSC.c1, Osteogenic.c1), some of these DE genes are shared across clusters, while some are specific to particular clusters (**Figure S3.1B**). Overall, there is a general decrease in interspecific DE as differentiation progresses and conversely. We found similar interspecific DE results when using Cormotif [12]. Specifically, we found that 5 correlation motifs best fit our pseudobulk data (**Figure S3.5D**). We detected 5,168 interspecific DE genes in iPSC.c1, 2,771 in iPSC.c2, 1,295 in iPSC.c1, 4,535 in MSC.c1, 3,585 in Osteogenic.c1, and 3,507 in Osteogenic.c2 with a posterior probability $> 0.65$ (**Figure S3.5E**), and several these DE genes are shared across clusters (**Figure S3.5F**).

Functional enrichments were examined (**Figure S3.1D-E, Figure S5.1B, Figure S5.3E, Figure S5.4E, Table S12, Table S13**). As expected, previously identified interspecific DE genes in iPSCs [13,14] were enriched among genes we identified as interspecific DE in iPSC.c1 (standard and Cormotif) but, for the most part, are not enriched among genes we identified as interspecific DE in other clusters (with several exceptions in the Cormotif tests). As a second check, previously identified interspecific DE genes in alternative cell types and tissues (non-pluripotent, non-mesenchymal, and non-osteogenic) [13,15] were, for the most, part not enriched among our DE genes (with some exceptions in both standard tests and Cormotif tests). On the other hand, previously identified interspecific DE genes in iPSC-derived CNCCs [14] are enriched among genes we identified as interspecific DE in MSC.c1 (Cormotif), in Osteogenic.c1 (standard), and across MSC.c1 and Osteogenic.c1 (standard and Cormotif). Additionally, previously identified interspecific DMR-associated genes identified in bone tissues [19] are enriched among genes we identified as interspecific DE in MSC.c1 (standard). Finally, regarding skeletal phenotype-related genes, almost no enrichments are present, except for genes associated with GWAS hits for height [20] which are enriched among genes we identified as interspecific DE in iPSC.c1 (Cormotif) and in MSC.c1 (Cormotif).

Lastly, we used Cormotif [12] to test for DE genes across stages of differentiation and found that 4 correlation motifs best fit our pseudobulk data when performing pairwise tests between subsequent clusters separately for each species (**Figure S3.1C**). These motifs are highly conserved across species, with about 85% of DE genes between clusters conserved across species. Of the remaining DE genes, 22 DE genes detected between iPSC.c1 and MSC.c1 differ between species, and 265 DE genes detected between MSC.c1 and Osteogenic.c1 differ between species.

**Identifying DE across an alternative general cell classification system (ad hoc assignments)**
Within each general ad hoc classification, we discovered hundreds of DE genes between species. Specifically, standard DE analyses of pseudobulk data detected 2,128 interspecific DE genes in iPSCs, 963 in MSCs, and 613 in osteogenic cells with an FDR < 0.01 (**Figure S3.2A**). Some of these DE genes are shared across ad hoc assignments, while some are specific to particular assignment (**Figure S3.2B**). Overall, there is a general decrease in interspecific DE as differentiation progresses. We found similar interspecific DE results when using Cormotif [12]. Specifically, we found that 2 correlation motifs best fit our pseudobulk data (**Figure S3.5G**). We detected 6,629 interspecific DE genes in iPSCs, 4,567 in MSCs, and 4,553 in osteogenic cells with a posterior probability > 0.65 (**Figure S3.5H**), and several these DE genes are shared across ad hoc assignments (**Figure S3.5I**).

Functional enrichments were examined (**Figure S3.2D-E, Figure S5.1C, Figure S5.3E, Figure S5.4E, Table S12, Table S13**). As expected, previously identified interspecific DE genes in iPSCs [13,14] were enriched among genes we identified as interspecific DE in iPSCs (standard and Cormotif) but, for the most part, are not enriched among genes we identified as interspecific DE in other ad hoc assignments (with some exceptions in the Cormotif tests). As a second check, previously identified interspecific DE genes in alternative cell types and tissues (non-pluripotent, non-mesenchymal, and non-osteogenic) [13,15] were, for the most part, not enriched among our DE genes (with some exceptions in both standard tests and Cormotif tests). On the other hand, previously identified interspecific DE genes in iPSC-derived CNCCs [14] are enriched among genes we identified as interspecific DE across MSCs and Osteogenic cells (standard and Cormotif). Additionally, previously identified interspecific DMR-associated genes identified in bone tissues [19] are enriched among genes we identified as interspecific DE in MSCs (standard) and across MSCs and Osteogenic cells (Cormotif). Finally, regarding skeletal phenotype-related genes, almost no enrichments are present, except for genes associated with GWAS hits for height [20] which are enriched among genes we identified as interspecific DE in iPSCs (Cormotif).

Lastly, we used Cormotif [12] to test for DE genes across stages of differentiation and found that 2 correlation motifs best fit our pseudobulk data when performing pairwise tests between subsequent ad hoc assignments separately for each species (**Figure S3.2C**). These motifs are not well conserved across species, though, with only 29% of DE genes between ad hoc assignments conserved across species. Of the remaining DE genes, 453 DE genes detected between iPSCs and MSCs differ between species, and 262 DE genes detected between MSCs and osteogenic cells differ between species.

**Comparing methods of assigning osteogenic cell annotations**
In addition to examining DE across general stages of differentiation, we established our comparative skeletal cell culture model as a more accessible way of specifically studying gene regulation within skeletal cell types. Using a broad grouping of osteogenic (Time 2) cells provided some insights, but increased cell heterogeneity at this time point as suggested by increased gene expression variance (**Figure 3E, Figure S3.7**) is a barrier. Thus, as described above, we tested several cell classification methods aimed at disentangling the approximate location of cells along the course of osteogenesis.

Using a flexible ad hoc assignment method that considers 4 candidate genes [10] as described above, we detected 2,004 preosteoblasts, 3,878 osteoblasts, 3,019 embedding osteoblasts, 4,473 mineralizing osteoblasts, and 2,443 maturing osteocytes (**Figure S4.1A**). Fewer human cells are assigned to each of these stages than chimpanzee cells (**Figure S4.1B**), but this is primarily because fewer human cells were recovered from the osteogenic cell (Time 2) collection (**Figure 2B**). Interestingly, although humans and chimpanzees have similar numbers of preosteoblasts, the distribution of cell counts along other stages of osteogenesis differs between species, with chimpanzees having an accumulation of cells at intermediate stages of osteogenesis (osteoblasts, embedding osteoblasts, and mineralizing osteoblasts) and humans having a decreased number of intermediate stages (embedding osteoblasts) but an increase in late stage cells (mineralizing osteoblasts and maturing osteocytes) (**Figure S4.1B**). These cell count distributions reflect the same pattern that we observed in the more traditional validation results, with chimpanzees on average producing less calcium deposits than humans (**Figure S1.2**), which indicates a larger proportion of osteogenic precursor cells in chimpanzees than humans. Moreover, it suggests that perhaps human cells can transition more quickly between intermediate and later stage osteogenic cells than chimpanzee cells. Finally, because these cells were assigned to a specific stage of osteogenesis based on their expression of known marker genes (**Figure 4B**), they retain these characteristic cell-specific expression patterns (**Figure S4.1C**) which allow for clear biological interpretations.

On the other hand, using an unsupervised clustering with a resolution of 0.50 as described above, we identified 4 osteogenic clusters (Osteogenic.c1, Osteogenic.c2, Osteogenic.c3, Osteogenic.c4) (**Figure S4.1D**). Similar to the osteogenic ad hoc assignment above, fewer human cells are assigned to each osteogenic cluster than chimpanzee cells (**Figure S4.1E**), but this is due to cell collection differences (**Figure 2B**). Conversely, within each species, similar proportions of cells are assigned to each cluster (**Figure S4.1E**). Although cells within these clusters display distinct patterns of candidate maker gene expression patterns (**Figure S4.1E**), these patterns are not always identical across species, nor do they clarify the stage at which cells are along the course of osteogenesis. Thus, because assigning cells using an ad hoc method produced more biologically meaningful groups of cells that are more consist with standard validation methods than using clustering methods, we proceeded with this ad hoc assignment scheme in the manuscript.

**Testing for concordance and functional enrichment of interspecific DE genes across stages of osteogenesis**

In order to ascertain the biological functions of interspecific DE genes identified across stages of osteogenesis (both via standard and Cormotif analyses), we tested for the enrichment of relevant external gene datasets (**Figure 4E-F, Figure S5.1D**) and GO categories (**Figure S5.3D, Figure S5.4D, Table S12, Table S13**). Resulting trends for the standard DE tests are supplied here, while details for the Cormotif DE tests are discussed in the manuscript.

As expected, previously identified inter-species DE genes in iPSCs [13,14] and in alternative cell types and tissues non-pluripotent, non-mesenchymal, and non-osteogenic) [13,15] are not enriched among genes we identified as interspecific DE in this study. Two exceptions are that previously identified interspecific DE genes in kidney tissue [15] are enriched among genes we identified as interspecific DE in osteoblasts and genes we identified as interspecific DE in embedding osteoblasts. Conversely, although previously identified interspecific DMR-associated genes identified in bone tissues [19] are not enriched among genes we identified as interspecific DE in this study, previously identified interspecific DE genes in CNCCs [14] are enriched among genes we identified as interspecific DE in preosteoblasts and genes we identified as interspecific DE in osteoblasts.

We found no overlap of skeletal trait and disease-related genes among interspecific DE genes. Specifically, we found no significant enrichment of hip geometry [17], bone area [18], height [20], osteoporosis [21], or osteoarthritis [22] related loci among interspecific DE genes identified in this study. Because there are very few genes associated with GWAS hits for hip geometry [17] and bone area [18] that also overlap with genes tested in this study, we also examined overlapping genes individually. However, we did not identify any genes that overlap. Finally, we did not find relevant GO category enrichment for most of our interspecific DE genes, with the exception of interspecific DE genes unique to preosteoblasts which are enriched in functions essential for skeletal tissue development and maintenance [23], such as Wnt signaling and tissue morphogenesis.

**Identifying DE across an alternative osteogenic cell classification system (unsupervised clusters)**

Within each unsupervised osteogenic cluster as described above, we discovered hundreds of DE genes between species. Specifically, standard DE analyses of pseudobulk data identified 643 interspecific DE genes in Osteogenic.c1, 1 in Osteogenic.c2, 428 in Osteogenic.c3, and 217 in Osteogenic.c4 with an FDR < 0.01 (**Figure S4.2B**). Some of these DE genes are shared across osteogenic clusters, while some are specific to particular clusters (**Figure S4.2C**). We found similar interspecific DE results when using Cormotif [12], although this method suggests a greater amount of sharing between Clusters. Specifically, we found that 2 correlation motifs best fit our pseudobulk data (**Figure S3.5M**). We identified 3,847 interspecific DE genes in Osteogenic.c1, 91 in Osteogenic.c2, 3,846 in Osteogenic.c3, and 3,813 in

Osteogenic.c4 with a posterior probability > 0.65 (**Figure S3.5N**), and almost all of these DE genes are shared across clusters (**Figure S3.5O**).

Functional enrichments were examined (**Figure S4.2D-E, Figure S5.1E, Figure S5.3E, Figure S5.4E, Table S12, Table S13**). As expected, previously identified interspecific DE genes in iPSCs [13,14] and in alternative cell types and tissues (non-pluripotent, non-mesenchymal, and non-osteogenic) [13,15] are not enriched among genes we identified as interspecific DE in this study. Conversely, although previously identified interspecific DE genes in iPSC-derived CNCCs [14] are not enriched among genes we identified as interspecific DE in this study, previously identified interspecific DMR-associated genes identified in bone tissues [19] are enriched among genes we identified as interspecific DE across Osteogenic.c1, Osteogenic.c3, and Osteogenic.c4 (Cormotif). Despite these enrichments, though, there are no significant overlap of skeletal phenotype- and disease-related genes which complicates functional interpretations.

#### Supplemental Tables

The following are descriptions of supplemental tables included in the manuscript.

**Table S1.** Details of samples used in the study.

**Table S2.** Known batch information for samples used in the study. Additional descriptions for each column heading are provided in **Table S2 (labels)**.

**Table S3.** Cell culture details for the MSC samples used in the study.

**Table S4.** Cell culture details for all samples used in the study.

**Table S5.** Details and results of the flow cytometry analyses.

**Table S6.** Details and results of the Alizarin Red S analyses.

**Table S7.** Details and results of the Oil Red O analyses.

**Table S8.** Details and results of the qPCR analyses.

**Table S9.** Known batch information for 10X single-cell sequencing collections performed in the study. Additional descriptions for each column heading are provided in **Table S9 (labels)**.

**Table S10.** The numbers of filtered genes tested in standard and Cormotif DE analyses across cell classifications.

**Table S11.** Summary of the external gene sets compiled for functional enrichment tests.

**Table S12.** Significant enrichment of GO functional categories in Cormotif DE genes identified for given cell classifications and inter-species DE gene sets.

**Table S13.** Significant enrichment of GO functional categories in standard DE genes identified for given cell classifications and inter-species DE gene sets.

#### Supplemental Figures

##### Figure S1.1

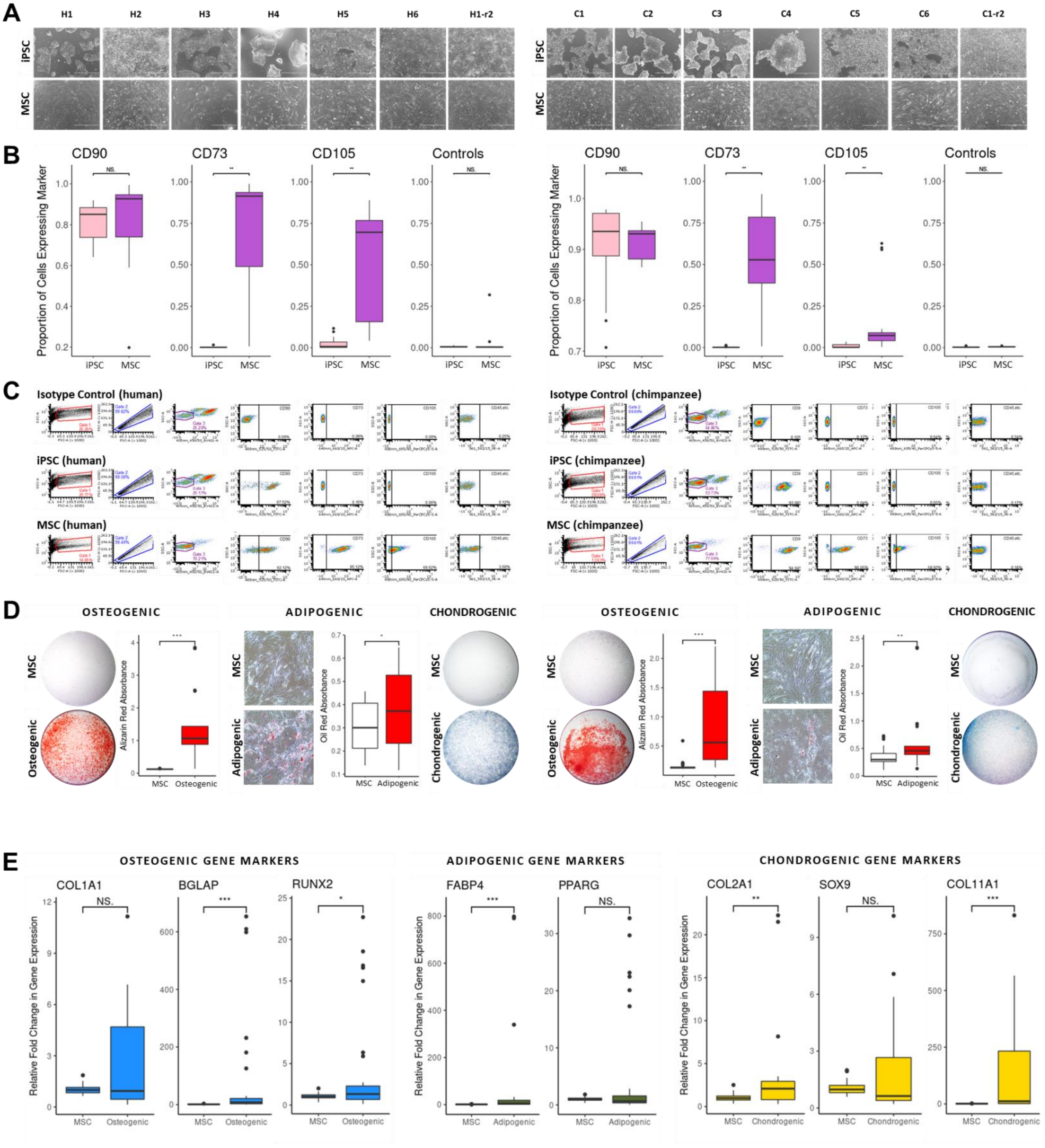

**Validation of MSC Differentiation.** (A) Phase contrast imaging at 4X of each cell line when pluripotent (Time 0, top row) and mesenchymal (Time 1, bottom row). (B) Box plots showing the proportion of cells expressing CD90, CD73, CD105, or control markers (CD45, CD34, CD14 or CD11b, CD19, and HLA-DR) for pluripotent cells (Time 0, pink) and mesenchymal cells (Time 1, purple) from all human biological and technical replicates (left set of plots) and all chimpanzee biological and technical replicates (right set of plots). Statistical significance was determined using one-sided Mann Whitney tests. (C) Representative flow cytometry analysis in one human cell line (left) and one chimpanzee cell line (right). Each set of plots displays results for the isotype control (top row), pluripotent cells (Time 0, second row), and mesenchymal cells (Time 1, bottom row). Plots in each row from left to right: (1) forward scatter area versus side scatter area of total cells, (2) forward scatter area versus forward scatter height of Gate 1 cells, (3) Zombie Violet fluorescence versus side scatter area of Gate 2 cells, (4) CD90 FITC fluorescence versus side scatter area of Gate 3 cells, (5) CD73 APC fluorescence versus side scatter area of Gate 3 cells, (6) CD105 PerCPCy5.5 fluorescence versus side scatter area of Gate 3 cells, (7) control marker (CD45, CD34, CD14 or CD11b, CD19, and HLA-DR) PE fluorescence versus side scatter area of Gate 3 cells. (D) Histological validation of MSC differentiation potential in human cell lines (left set of plots) and chimpanzee cell lines (right set of plots). Plots in each set from left to right: (1) images of mesenchymal cells (Time 1) and osteogenic cells (Time 2) after Alizarin Red staining (image zoomed out to display the entire cell culture well), (2) box plot of absorbance values of extracted Alizarin Red stain from all biological and technical replicates from a given species, (3) phase contrast image at 4X magnification of mesenchymal cells (Time 1) and adipogenic cells after Oil Red staining, (4) box plot of absorbance values of extracted Oil Red stain from all biological and technical replicates from a given species, (5) images of mesenchymal cells (Time 1) and chondrogenic cells after Alcian Blue staining (image zoomed out to display the entire cell culture well). (E) Box plots showing the relative quantification levels calculated using the $E^{-\Delta\Delta C_T}$ method of osteogenic (*COL1A1, BGLAP, RUNX2*), adipogenic (*FABP4, PPARG*), and chondrogenic (*COL2A1, SOX9, COL11A1*) candidate genes in mesenchymal cells (Time 1) and osteogenic cells (Time 2) in all samples. Statistical significance was determined using one-sided t-tests on ΔΔCt values.

Box plots: middle line marks the median, box outlines the first and third quartiles, whiskers extend to 1.5 times the interquartile range

Significance: NS. p>0.05, * p<0.05, ** p<0.01, *** p <0.001

**Figure S1.2**

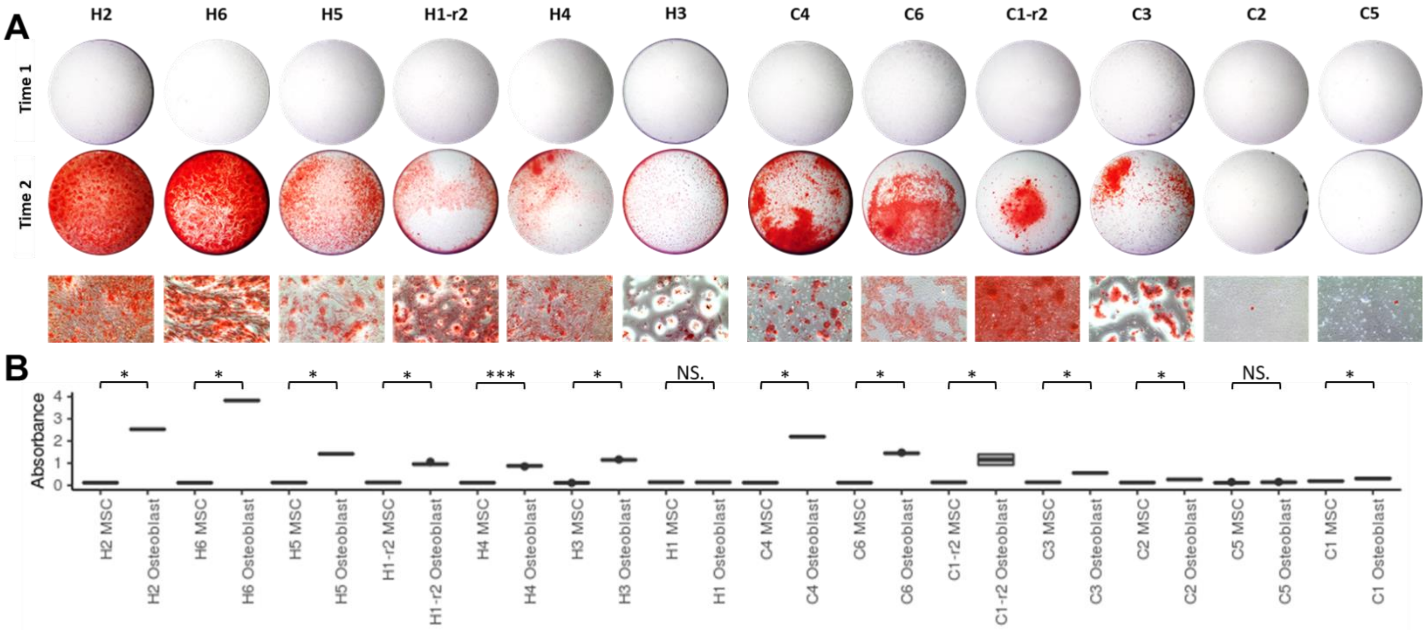

**Validation of Osteogenic Differentiation.** (A) Imaging of Alizarin Red stain in cell types from each cell line. Images in each row from top to bottom: (1) stained mesenchymal cells (Time 1) zoomed out to display the entire cell culture well, (2) stained osteogenic cells (Time 2) zoomed out to display the entire cell culture well, (3) phase contrast imaging at 4X of stained osteogenic cells (Time 2). (B) Box plots showing the absorbance values of extracted Alizarin Red stain from all biological and technical replicates. Statistical significance was determined using one-sided Mann Whitney tests.

Box plots: middle line marks the median, box outlines the first and third quartiles, whiskers extend to 1.5 times the interquartile range

Significance: NS. p>0.05, * p<0.05, ** p<0.01, *** p <0.001

**Figure S2.1**

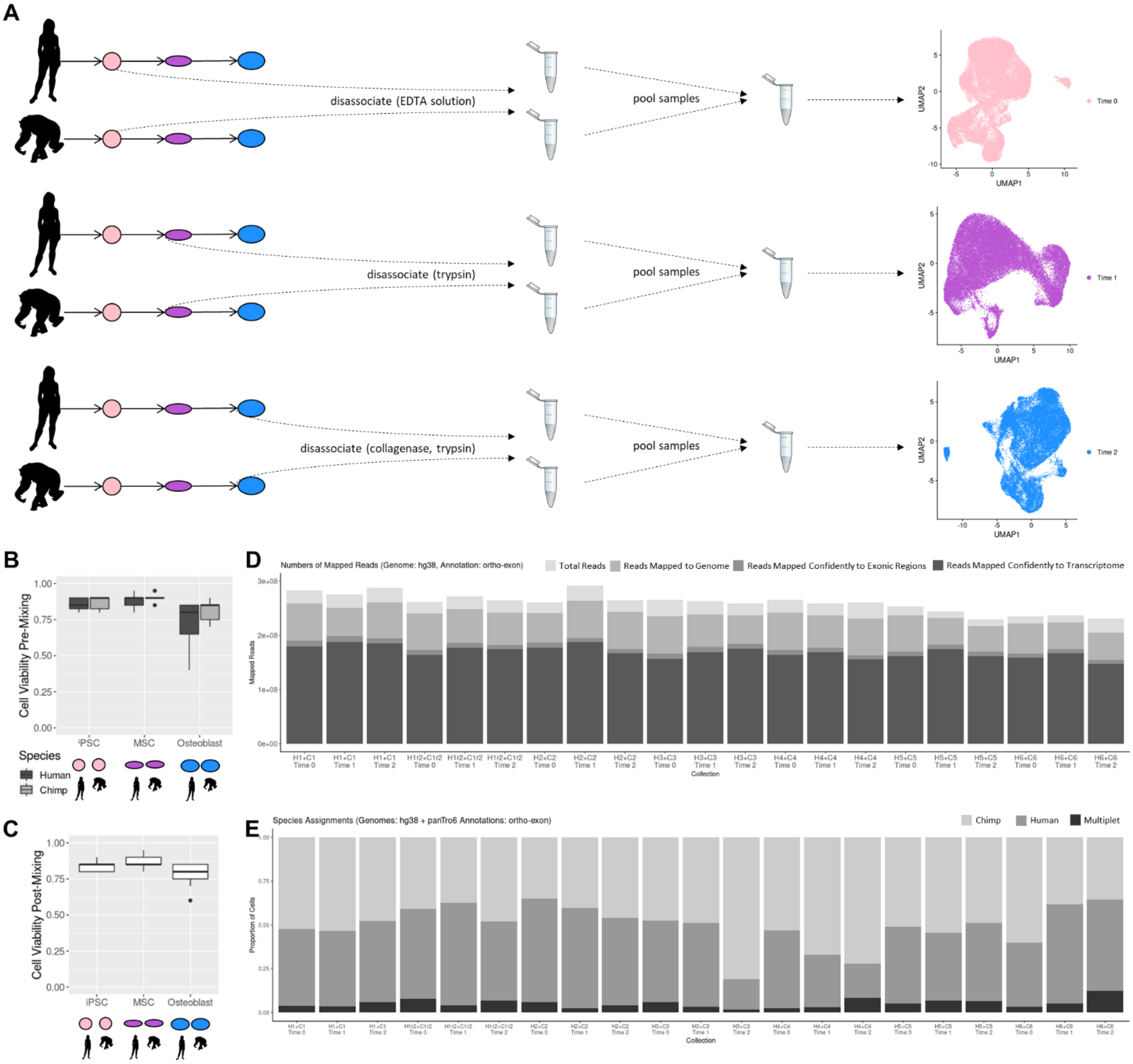

**10X Single-Cell Collections.** (A) Schematic of the dissociation and pooling protocol for 10X single-cell collections. (B) Box plots show cell viabilities for each species and cell type after disassociation. (C) Box plots show cell viabilities for pooled samples of human and chimpanzee cells after mixing. (D) Proportion of mapped scRNA-seq reads. (E) Proportions of species assignments for each 10X collection using Cell Ranger.

Box plots: middle line marks the median, box outlines the first and third quartiles, whiskers extend to 1.5 times the interquartile range

Silhouette images were adapted from http://phylopic.org/.

**Figure S2.2**

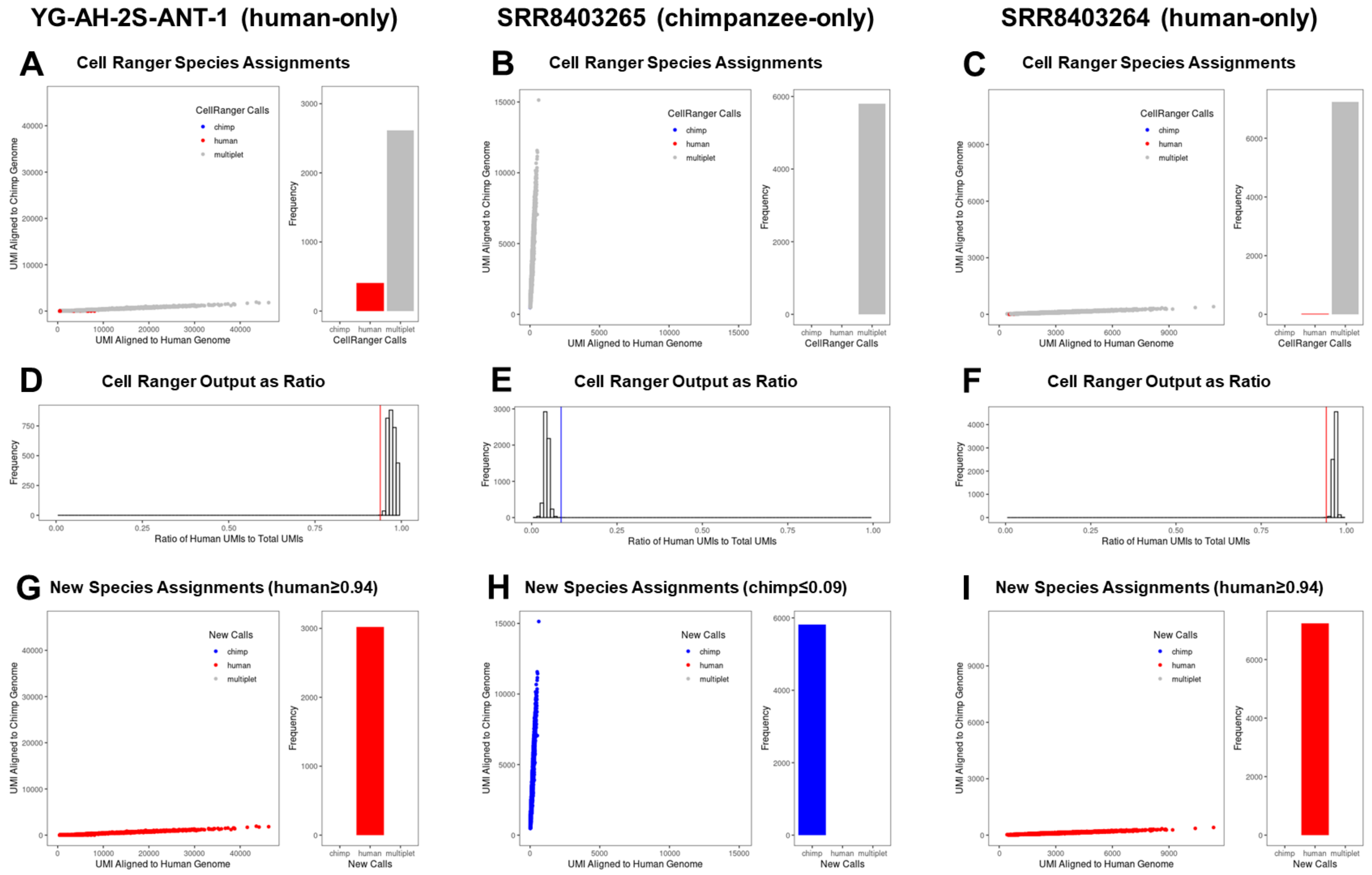

**Testing Cell Ranger Species Assignments.** Results of running Cell Ranger on one dataset that contained only human cells [YG-AH-2S-ANT-1] (A,D,G), one dataset that contained only chimpanzee cells [SRR8403265] (B,E,H), and one dataset that contained only human cells [SRR8403264] (C,F,I). Top plots show species assignments for each cell when using the standard Cell Ranger pipeline (A,B,C). Plots in each set from left to right: (1) UMI counts aligned to the human genome vs. UMI counts aligned to the chimpanzee genome with cells colored by assignment, (2) number of cells called as human, chimpanzee, or multiplet. Middle plot shows a histogram of the ratio of human-aligned UMI counts per cells vs. the total number of aligned UMI counts per cells (bars near 0 are likely chimpanzee cells and bars near 1 are likely human cells) (D,E,F). Bottom plots show an alternative species assignment using low cutoffs (~0.9 for humans and ~0.1 for chimps) based on the ratios shown in the histograms (G,H,I). Plots in each set from left to right: (1) UMI counts aligned to the human genome vs. UMI counts aligned to the chimpanzee genome with cells colored by assignment, (2) number of cells called as human, chimpanzee, or multiplet.

**Figure S2.3**

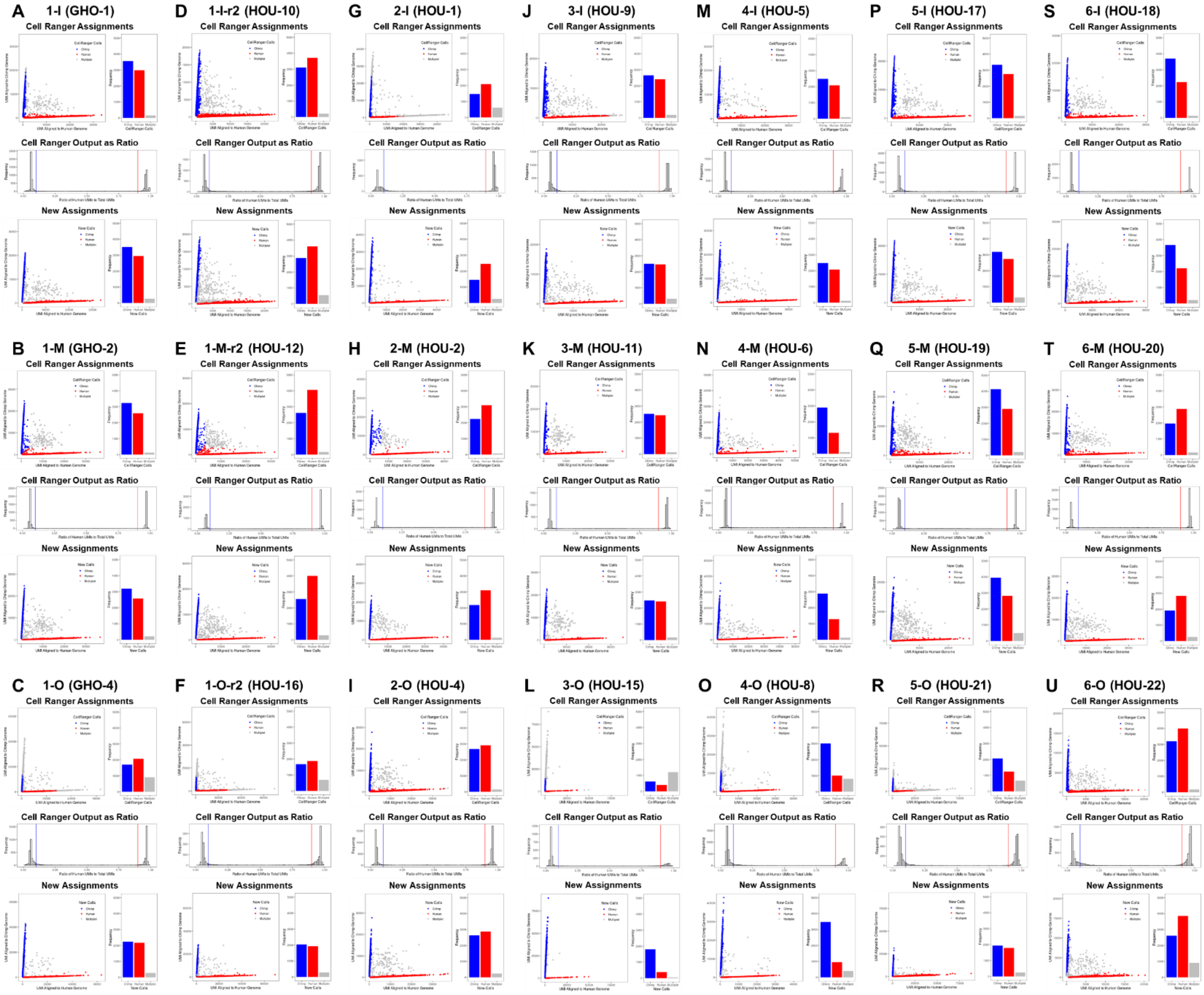

**Applying Modification to Cell Ranger Species Assignments.** Results of running Cell Ranger vs. our modification on all datasets in the study. Top plots show species assignments for each cell when using the standard Cell Ranger pipeline (A,B,C,J,M,P,S). Plots in each set from left to right: (1) UMI counts aligned to the human genome vs. UMI counts aligned to the chimpanzee genome with cells colored by assignment, (2) number of cells called as human, chimpanzee, or multiplet. Middle plot shows a histogram of the ratio of human-aligned UMI counts per cells vs. the total number of aligned UMI counts per cells (bars near 0 are likely chimpanzee cells and bars near 1 are likely human cells) (D,E,F,K,N,Q,T). Bottom plots show an alternative species assignment using low cutoffs (~0.9 for humans and ~0.1 for chimps) based on the ratios shown in the histograms (G,H,I,L,O,R,U). Plots in each set from left to right: (1) UMI counts aligned to the human genome vs. UMI counts aligned to the chimpanzee genome with cells colored by assignment, (2) number of cells called as human, chimpanzee, or multiplet.

**Figure S2.4**

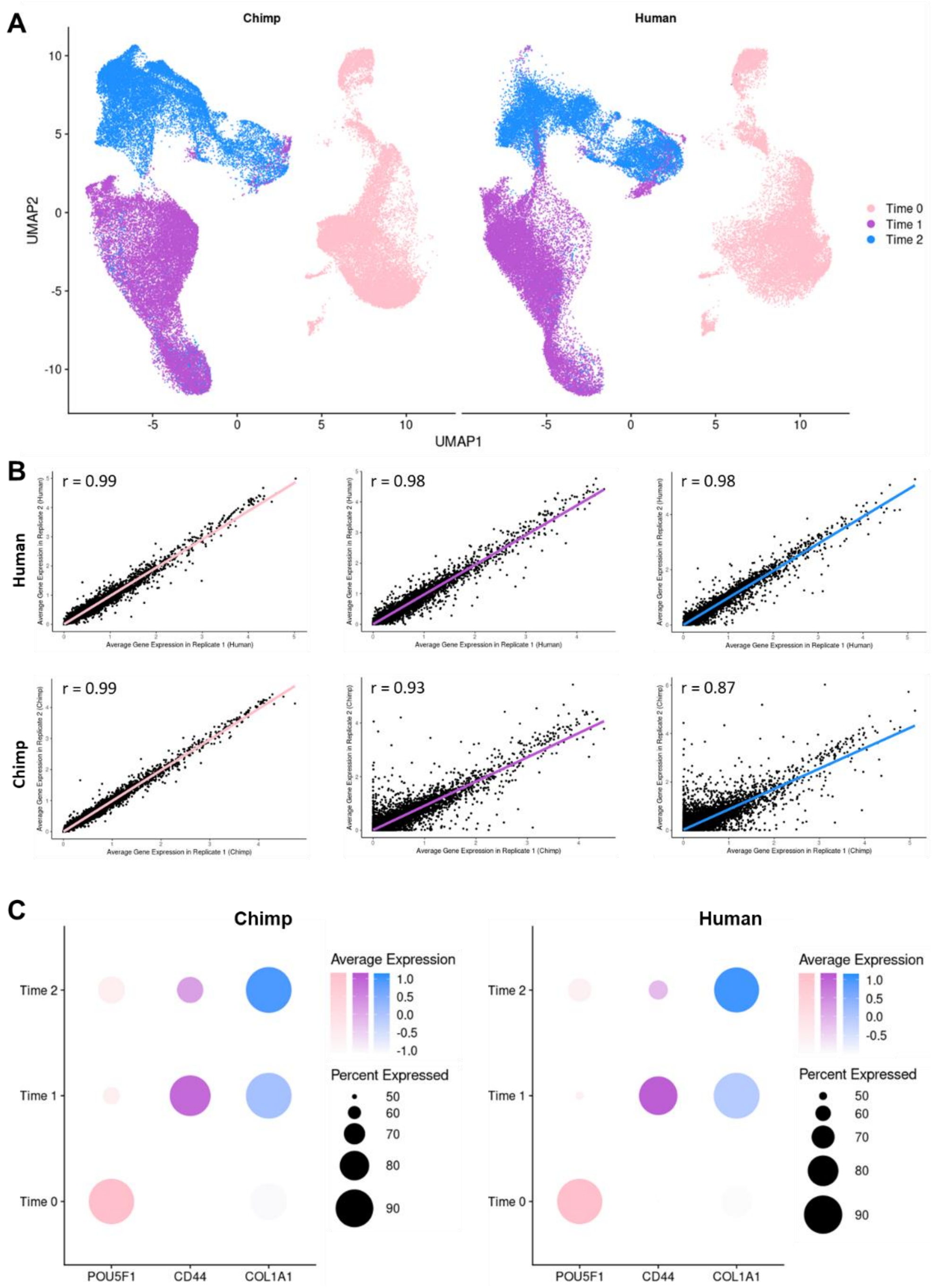

**Data Integration, Reproducibility, and Candidate Gene Expression.** (A) UMAP dimensional reduction plots of scRNA-seq data with cells labeled by the stage of differentiation at which they were collected and separated by species. (B) The correlation of average gene expression patterns between technical replicates separated by species. Plots from left to right: (1) correlation in pluripotent cells (Time 0), (2) correlation in mesenchymal cells (Time 1), (3) correlation in osteogenic cells (Time 2). (C) Dot plots depicting the scaled average expression (dot color intensity) and the proportion of cells expressing each gene (dot size) of candidate genes (x-axis) at each stage of differentiation (y-axis).

**Figure S2.5**

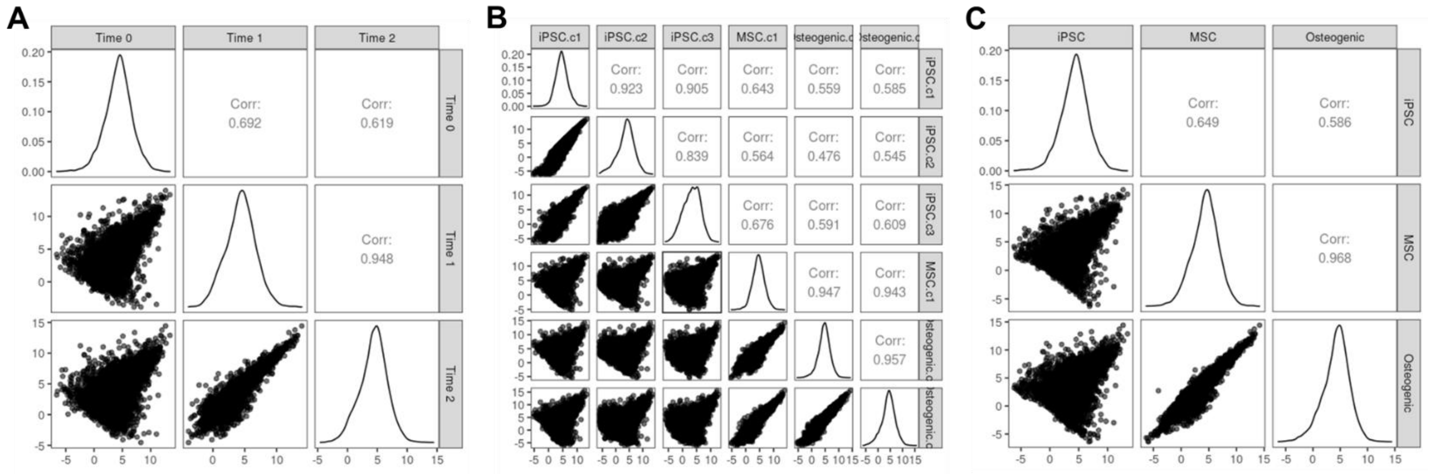

**Whole Transcriptome Correlations in General Cell Classifications.** (A) Pairwise correlations of pseudobulk counts for each gene between stages of differentiation. (B) Pairwise correlations of pseudobulk counts for each gene between unsupervised clusters at a resolution of 0.05. (C) Pairwise correlations of pseudobulk counts for each gene between general ad hoc assignments.

**Figure S2.6**

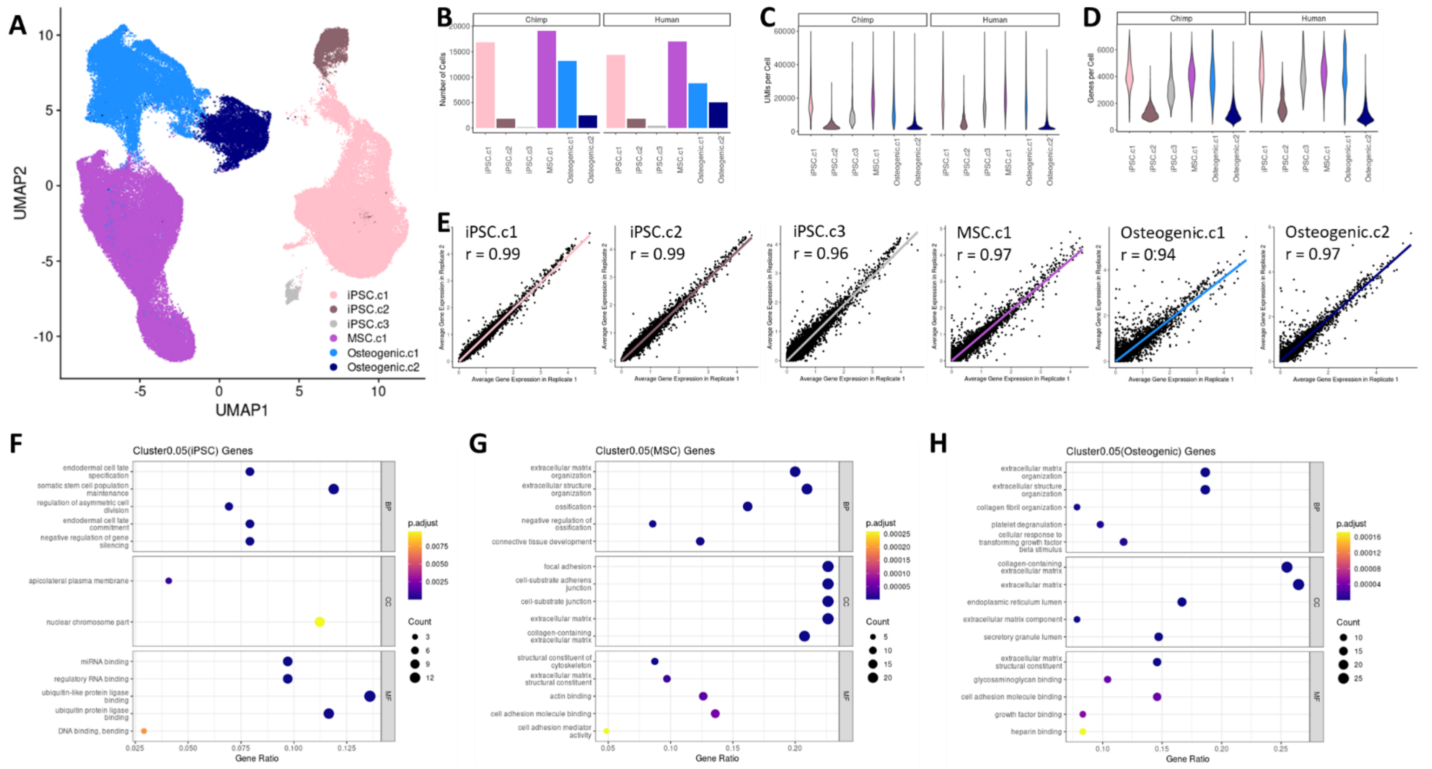

**Data QC in General Unsupervised Cluster Cell Annotations.** (A) UMAP dimensional reduction plot of scRNA-seq data with cells labeled by their assigned unsupervised cluster (resolution=0.05). (B) Bar plot depicting the number of chimpanzee and human cells assigned to each cluster. (C) Violin plots displaying the distribution of UMI counts per cell for each cluster. (D) Violin plots displaying the distribution of gene counts per cell for each cluster. (E) The correlation of average gene expression patterns between technical replicates collected in each cluster. (F-H) Enrichment of GO functional categories in marker genes for iPSC.c1 (F), MSC.c1 (G), and Osteogenic.c1 (H). The top 5 GO functions identified in biological processes (BP), cell components (CC), and molecular functions (MF) are displayed along with the adjusted p-value (p-adjust), the number of marker genes overlapping a GO function (Count), and the ratio of overlapping to non-overlapping marker genes for a given GO function (GeneRatio).

**Figure S2.7**

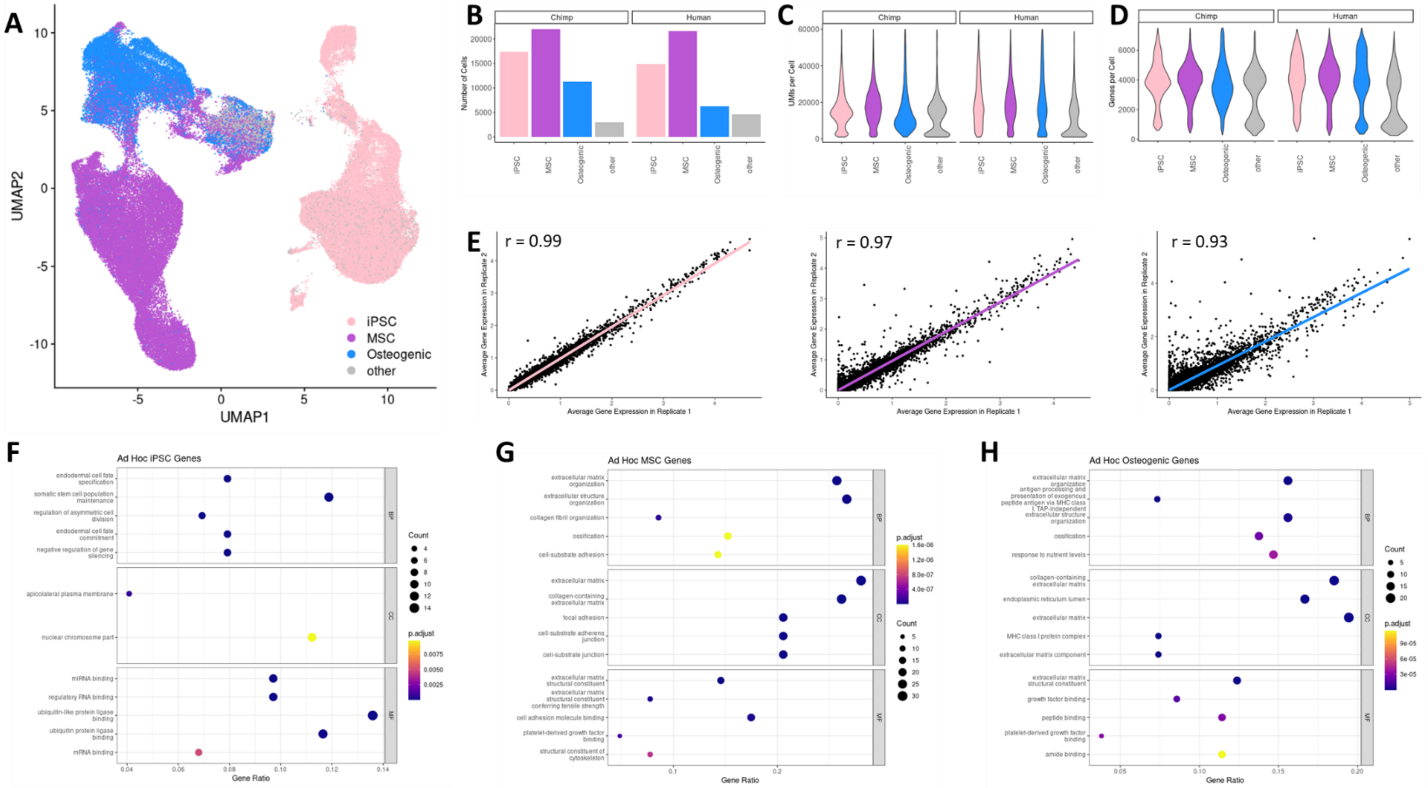

**Data QC in General Ad Hoc Assignment Cell Annotations.** (A) UMAP dimensional reduction plot of scRNA-seq data with cells labeled by their ad hoc assignment. (B) Bar plot depicting the number of chimpanzee and human cells in each ad hoc assignment. (C) Violin plots displaying the distribution of UMI counts per cell for each ad hoc assignment. (D) Violin plots displaying the distribution of gene counts per cell for each ad hoc assignment. (E) The correlation of average gene expression patterns between technical replicates collected in each ad hoc assignment. (F-H) Enrichment of GO functional categories in marker genes for iPSC.c1 (F), MSC.c1 (G), and Osteogenic.c1 (H). The top 5 GO functions identified in biological processes (BP), cell components (CC), and molecular functions (MF) are displayed along with the adjusted p-value (p-adjust), the number of marker genes overlapping a GO function (Count), and the ratio of overlapping to non-overlapping marker genes for a given GO function (GeneRatio).

**Figure S3.1**

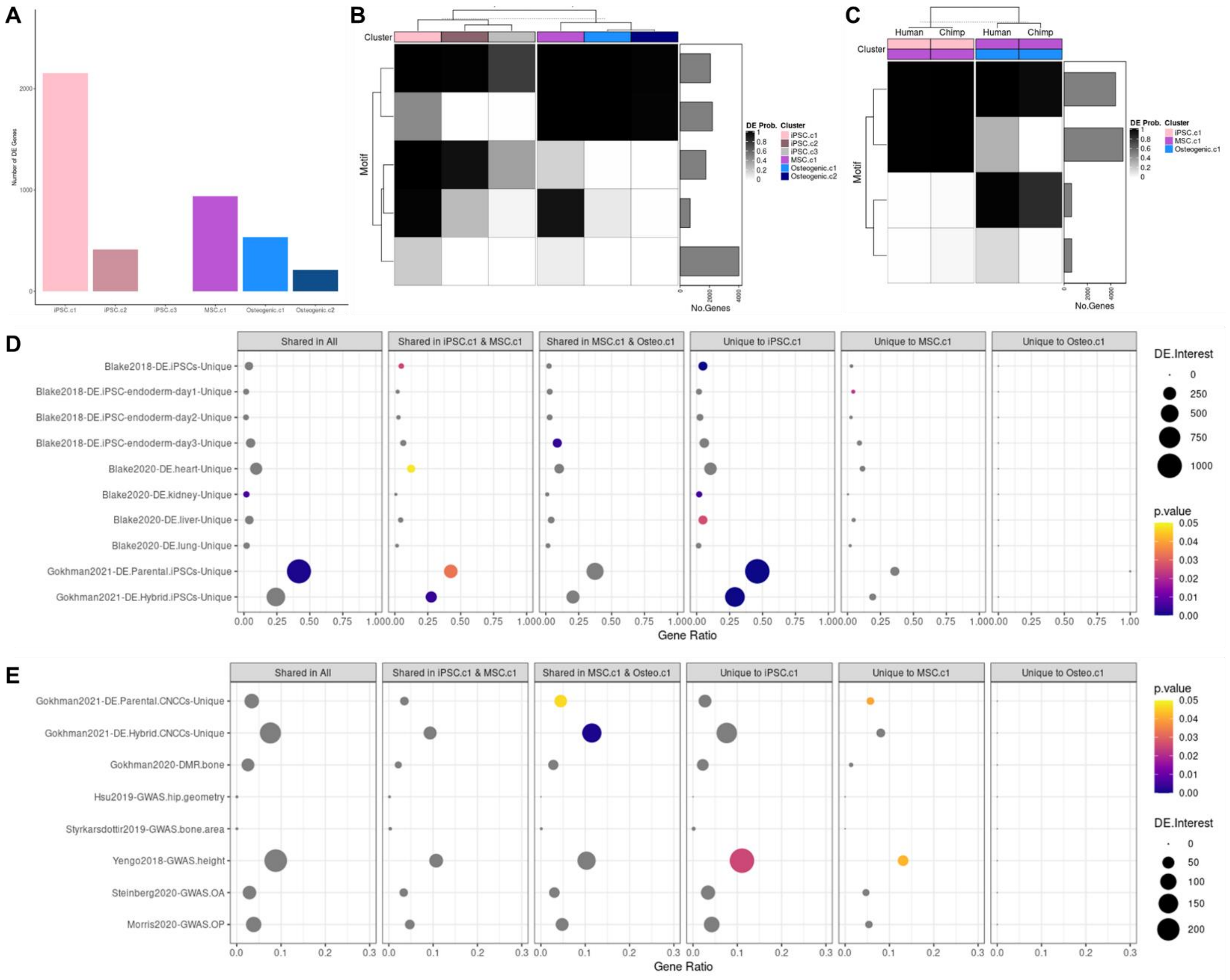

**Interspecific DE in General Unsupervised Cluster Cell Annotations.** (A) Bar plot showing the number of interspecific DE genes identified for each unsupervised cluster (resolution=0.05) using standard methods. (B-C) Correlation motifs based on the probability of differential expression between species for each cluster (B) and correlation motifs based on the probability of differential expression across clusters (iPSC.c1, MSC.c1, Osteogenic.c1) for each species (C) with the number of genes assigned to each motif shown in the bar plot on the right and the posterior probability that a gene is DE between two clusters in a given species shown by the shading of each box. (D-E) Enrichment of external DE gene sets among Cormotif interspecific DE genes identified for each cluster with the p-value (p.value), the number of DE genes overlapping an external gene set (DE.Interest), and the ratio of overlapping to non-overlapping DE genes for a given external gene set (GeneRatio) denoted.

**Figure S3.2**

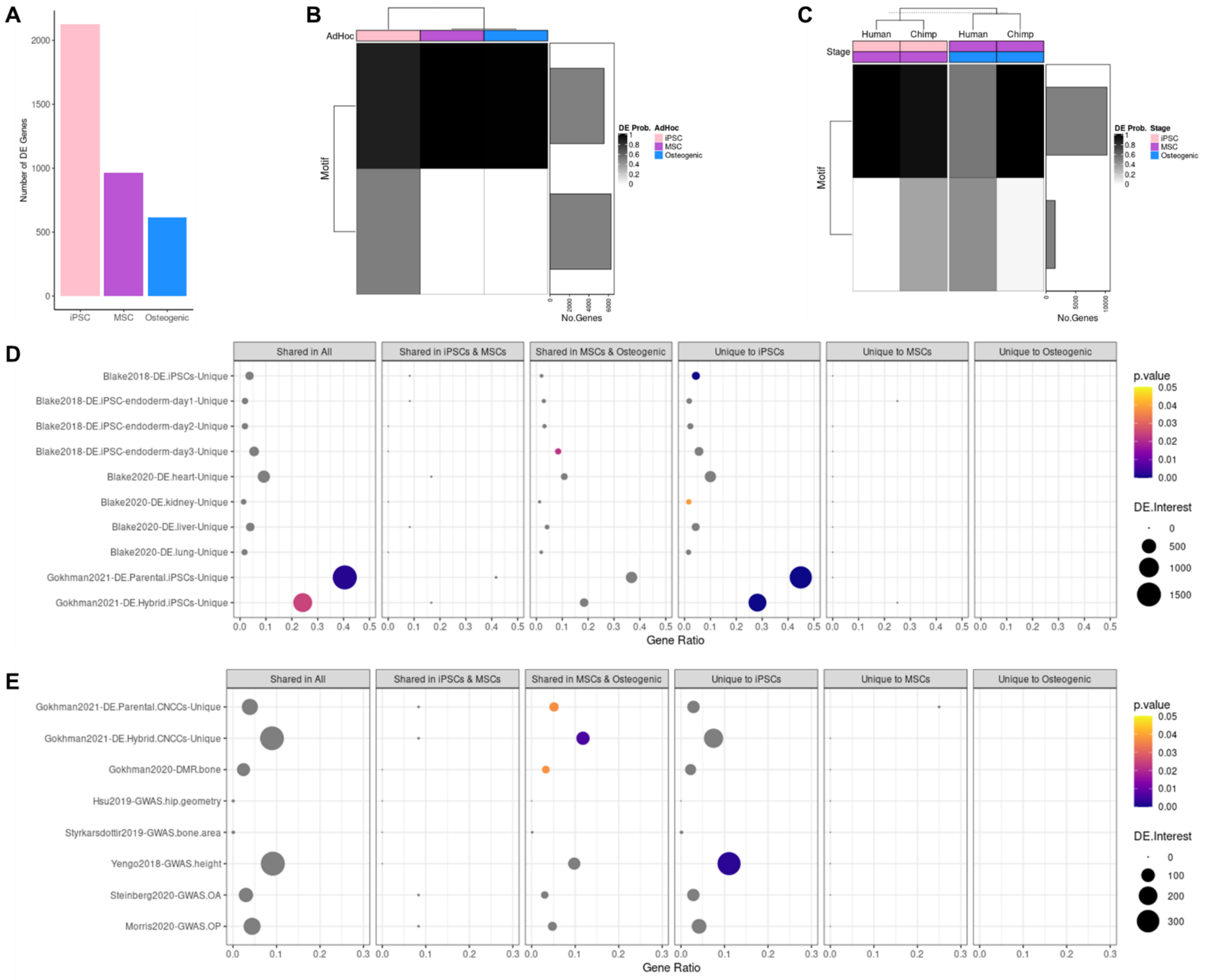

**Interspecific DE in General Ad Hoc Assignment Cell Annotations.** (A) Bar plot showing the number of standard interspecific DE genes identified for each ad hoc assignment. (B-C) Correlation motifs based on the probability of differential expression between species for each ad hoc assignment (B) and correlation motifs based on the probability of differential expression across ad hoc assignments for each species (C) with the number of genes assigned to each motif shown in the bar plot on the right and the posterior probability that a gene is DE between two clusters in a given species shown by the shading of each box. (D-E) Enrichment of external DE gene sets among Cormotif interspecific DE genes identified for each ad hoc assignment with the p-value (p.value), the number of DE genes overlapping an external gene set (DE.Interest), and the ratio of overlapping to non-overlapping DE genes for a given external gene set (GeneRatio) denoted.

#### Figure S3.3

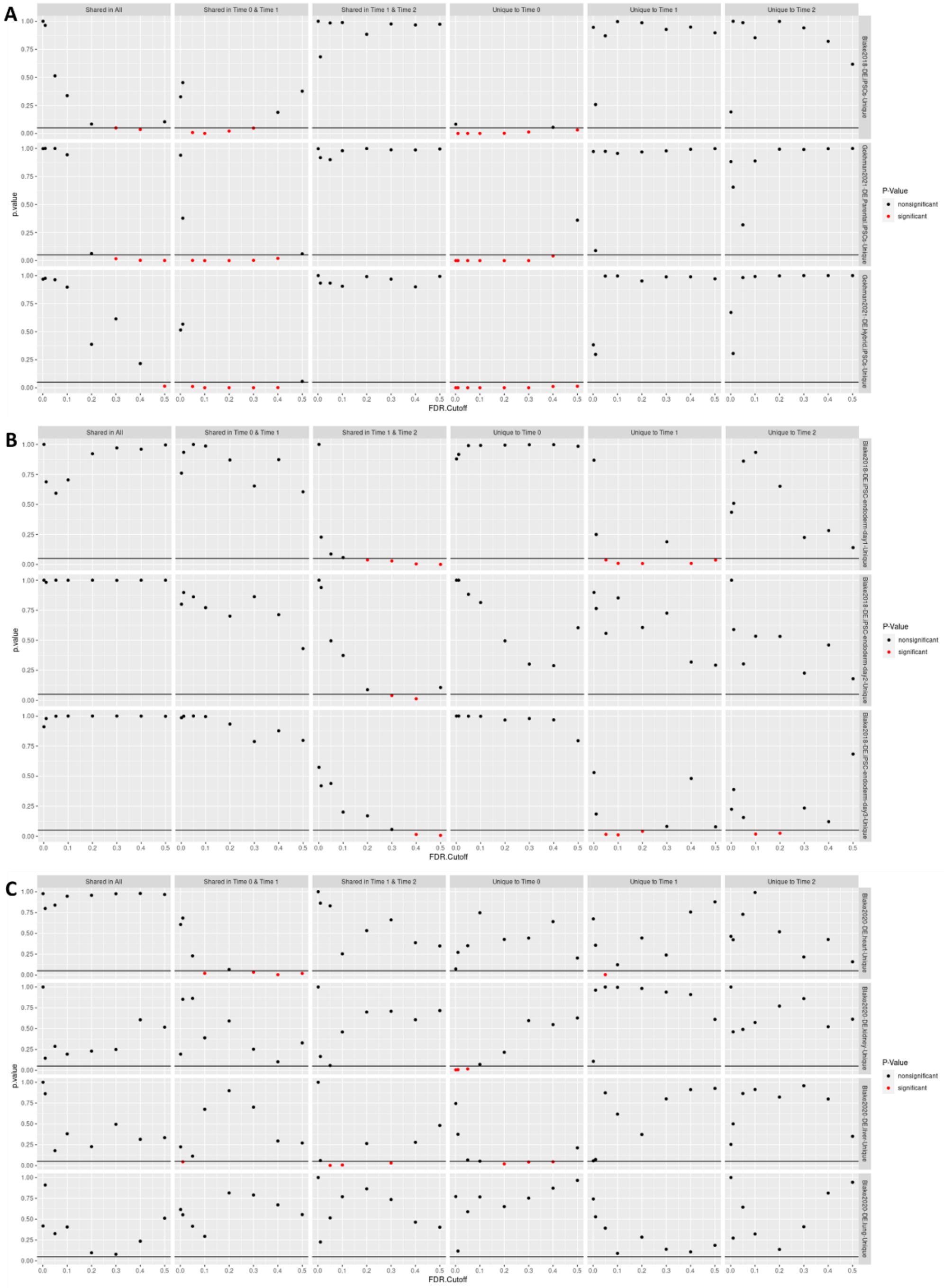

**Assessing FDR Thresholds for Calling Standard DE Genes.** Enrichment p-values (p.value) of external DE gene sets among interspecific DE genes identified across stages of differentiation using standard methods and a range of different FDR cutoffs (FDR.Cutoff). A p-value of 0.05 is denoted by a horizontal line on each plot, and significant enrichments (p-value < 0.05) are highlighted in red. External DE gene sets were chosen for validation purposes – (A) previously identified interspecific DE genes in iPSCs are expected to only be enriched in interspecific DE genes unique to pluripotent cells (Time 0), (B) previously identified interspecific DE genes in alternative cell types (non-pluripotent, non-mesenchymal, and non-osteogenic) are not expected to be enriched in any interspecific DE genes identified in this study, and (C) previously identified interspecific DE genes in alternative tissue types (non-pluripotent, non-mesenchymal, and non-osteogenic) are not expected to be enriched in any interspecific DE genes identified in this study.

**Figure S3.4**

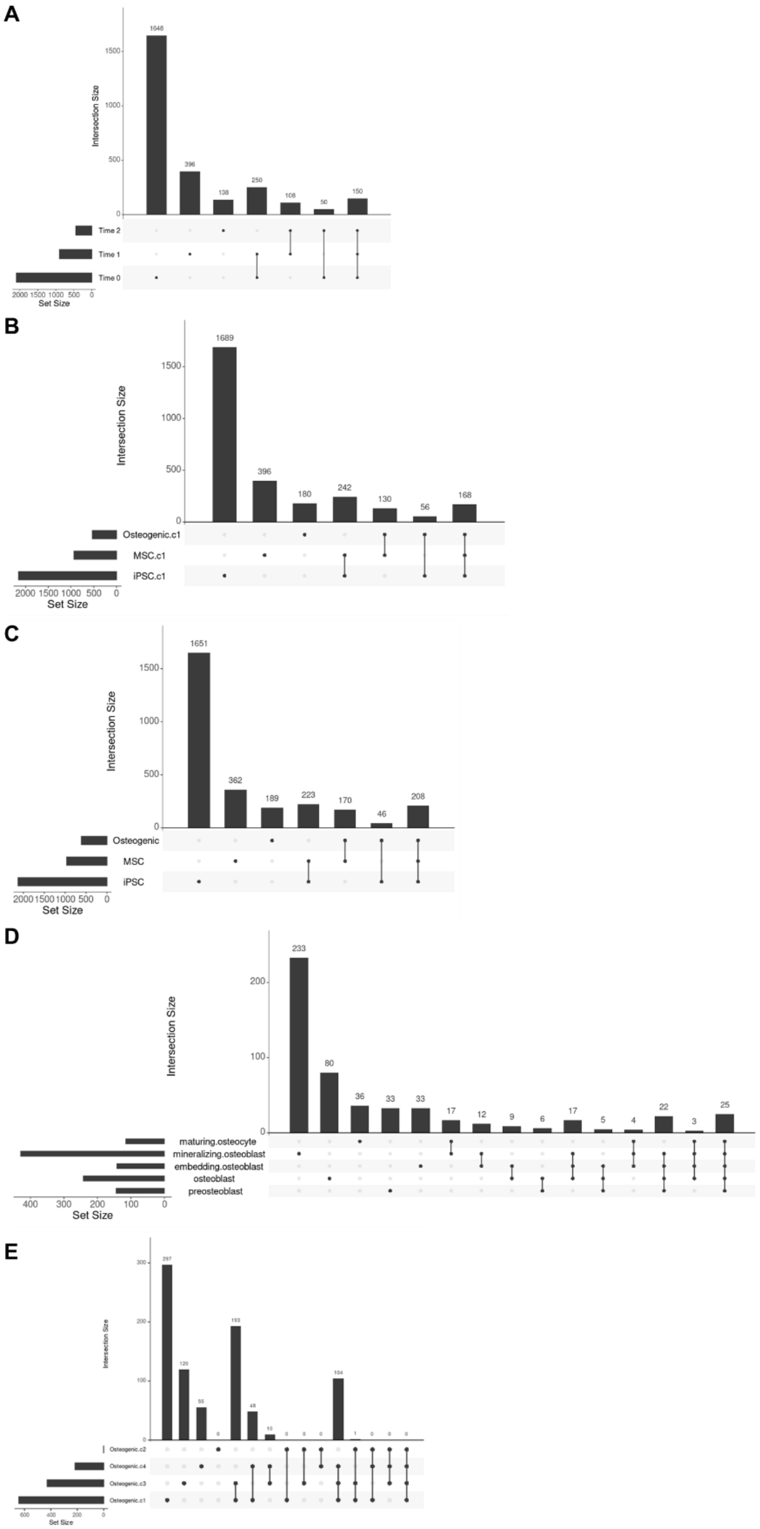

**Overlap of Standard Interspecific DE Genes.** (A) Details regarding the intersection of standard interspecific DE genes identified across stages of differentiation. (B) Details regarding the intersection of standard interspecific DE genes identified across general unsupervised clusters (resolution=0.05). (C) Details regarding the intersection of standard interspecific DE genes identified across general ad hoc assignments. (D) Details regarding the intersection of standard interspecific DE genes identified across osteogenic ad hoc assignments. (E) Details regarding the intersection of standard interspecific DE genes identified across osteogenic unsupervised clusters (resolution=0.50).

**Figure S3.5**

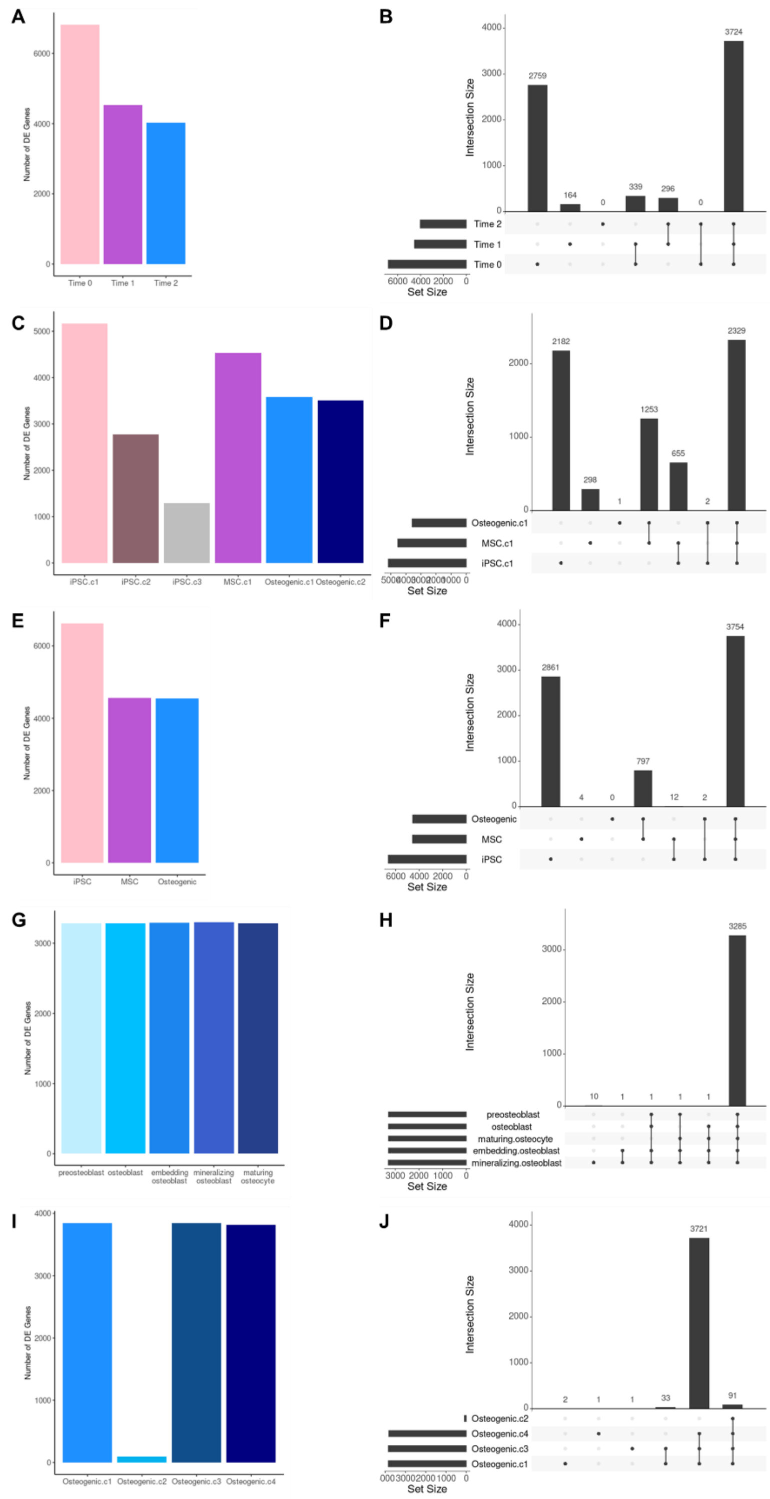

**Interspecific Cormotif DE Genes.** (A-B) Plots describing Cormotif interspecific DE genes identified across stages of differentiation. (C-D) Plots describing Cormotif interspecific DE genes identified across general unsupervised clusters (resolution=0.05). (E-F) Plots describing Cormotif interspecific DE genes identified across general ad hoc assignments. (G-H) Plots describing Cormotif interspecific DE genes identified across osteogenic ad hoc assignments. (I-J) Plots describing Cormotif interspecific DE genes identified across osteogenic unsupervised clusters (resolution=0.50). Plots within each set from left to right: (1) bar plot showing the number of interspecific DE genes identified using Cormotif for given cell classifications (A,C,E,G,I), (2) details regarding the intersection of DE genes for given cell classification (B,D,F,H,J).

**Figure S3.6**

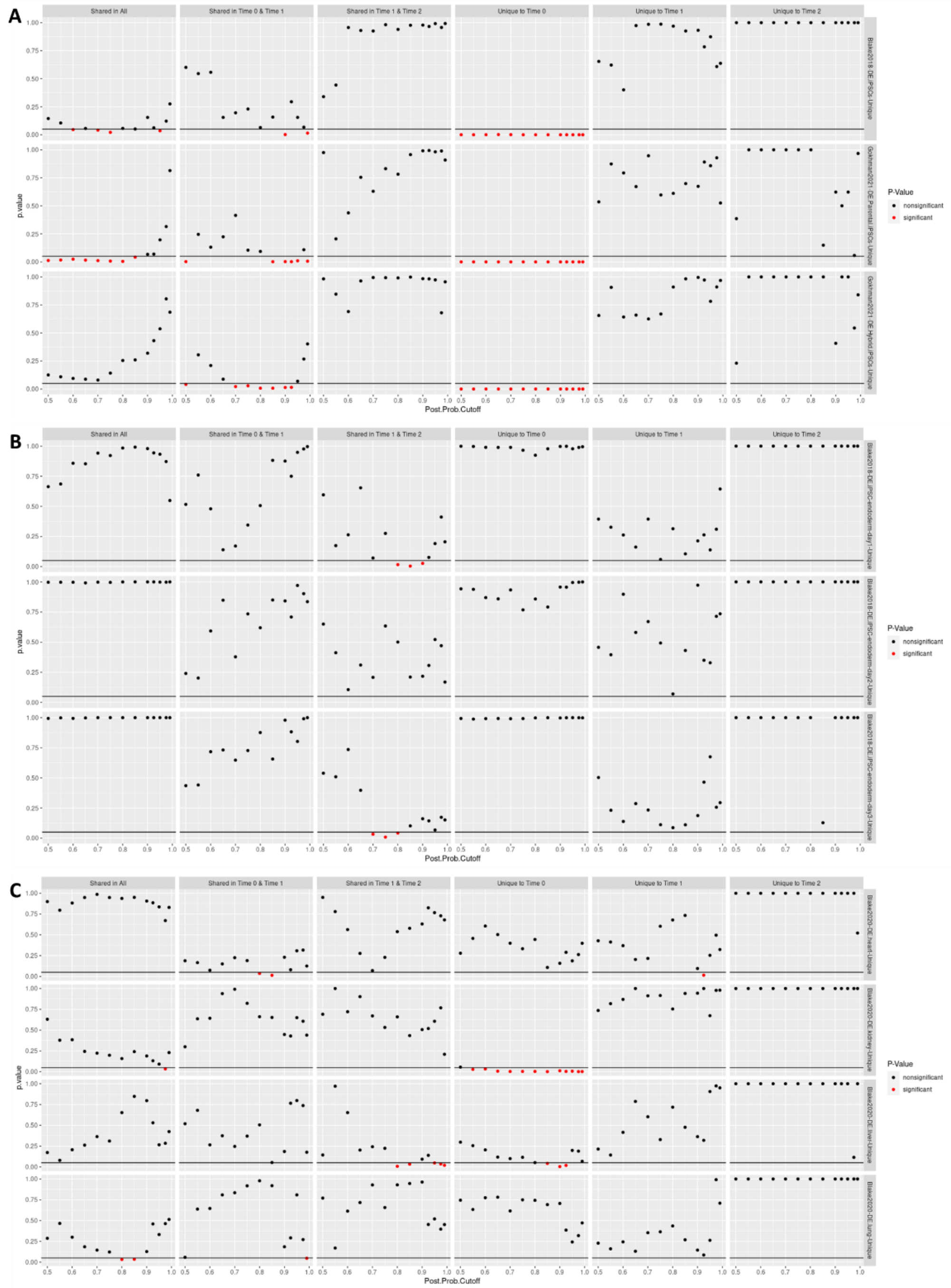

**Assessing Posterior Probability Thresholds for Calling Cormotif DE Genes.** Enrichment p-values (p.value) of external DE gene sets among Cormotif interspecific DE genes identified across stages of differentiation using a range of different posterior probability cutoffs (Post.Prob.Cutoff). A p-value of 0.05 is denoted by a horizontal line on each plot, and significant enrichments (p-value < 0.05) are highlighted in red. External DE gene sets were chosen for validation purposes – (A) previously identified interspecific DE genes in iPSCs are expected to only be enriched in interspecific DE genes unique to pluripotent cells (Time 0), (B) previously identified interspecific DE genes in alternative cell types (non-pluripotent, non-mesenchymal, and non-osteogenic) are not expected to be enriched in any interspecific DE genes identified in this study, and (C) previously identified interspecific DE genes in alternative tissue types (non-pluripotent, non-mesenchymal, and non-osteogenic) are not expected to be enriched in any interspecific DE genes identified in this study.

**Figure S3.7**

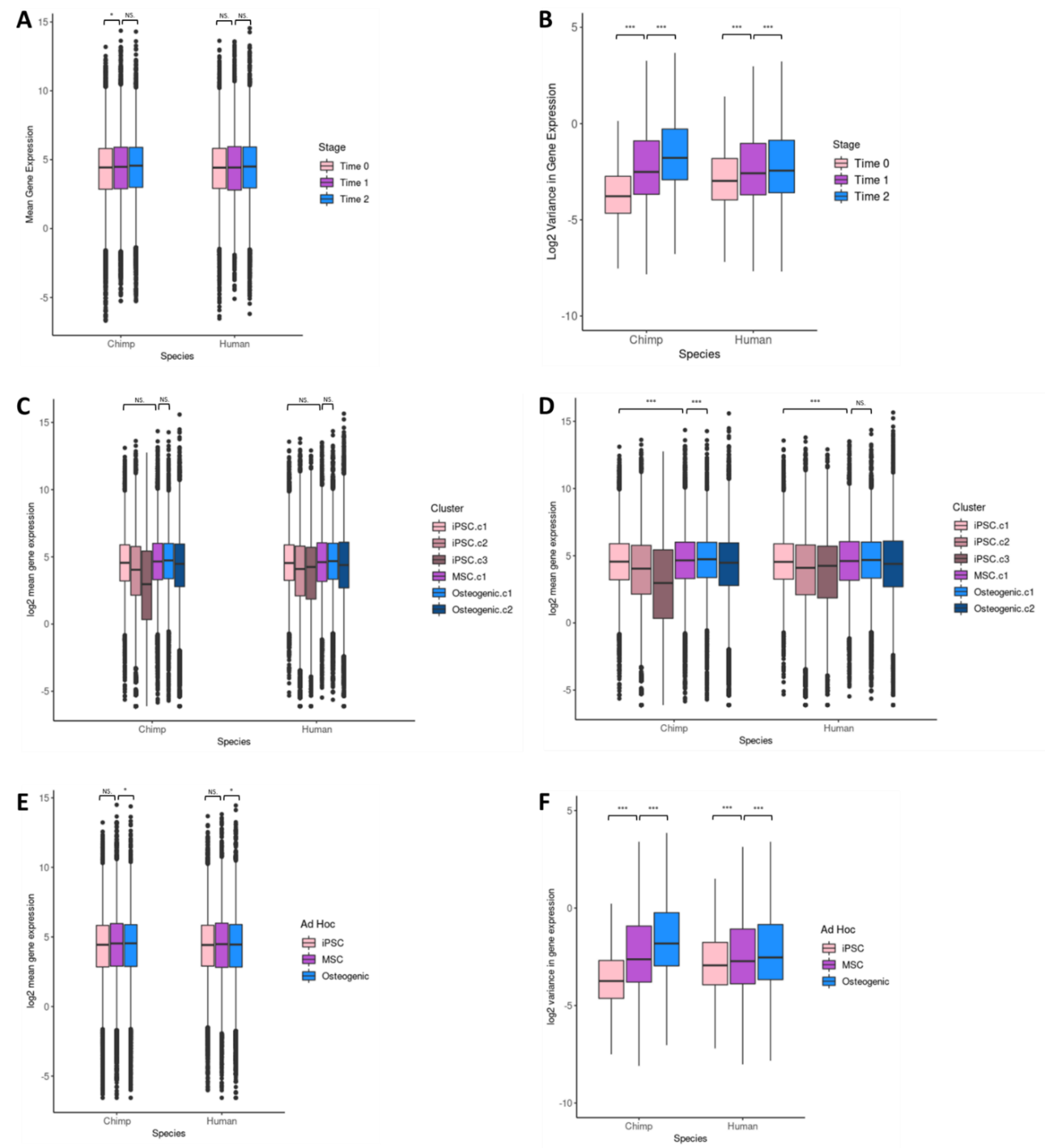

**Gene Expression Means and Variances.** (A,C,E) Box plots of the mean gene expression values for given cell classifications in each species. (B,D,F) Box plots of the log2 transformed gene expression variance values for given cell classifications in each species. Statistical significance was determined using two-sided t-tests. (A-B) Mean and variance values for stages of differentiation. (C-D) Mean and variance values for general unsupervised clusters (resolution=0.05). (E-F) Mean and variance values for general ad hoc assignments.

Box plots: middle line marks the median, box outlines the first and third quartiles, whiskers extend to 1.5 times the interquartile range

Significance: NS. p>0.05, * p<0.05, ** p<0.01, *** p <0.001

**Figure S4.1**

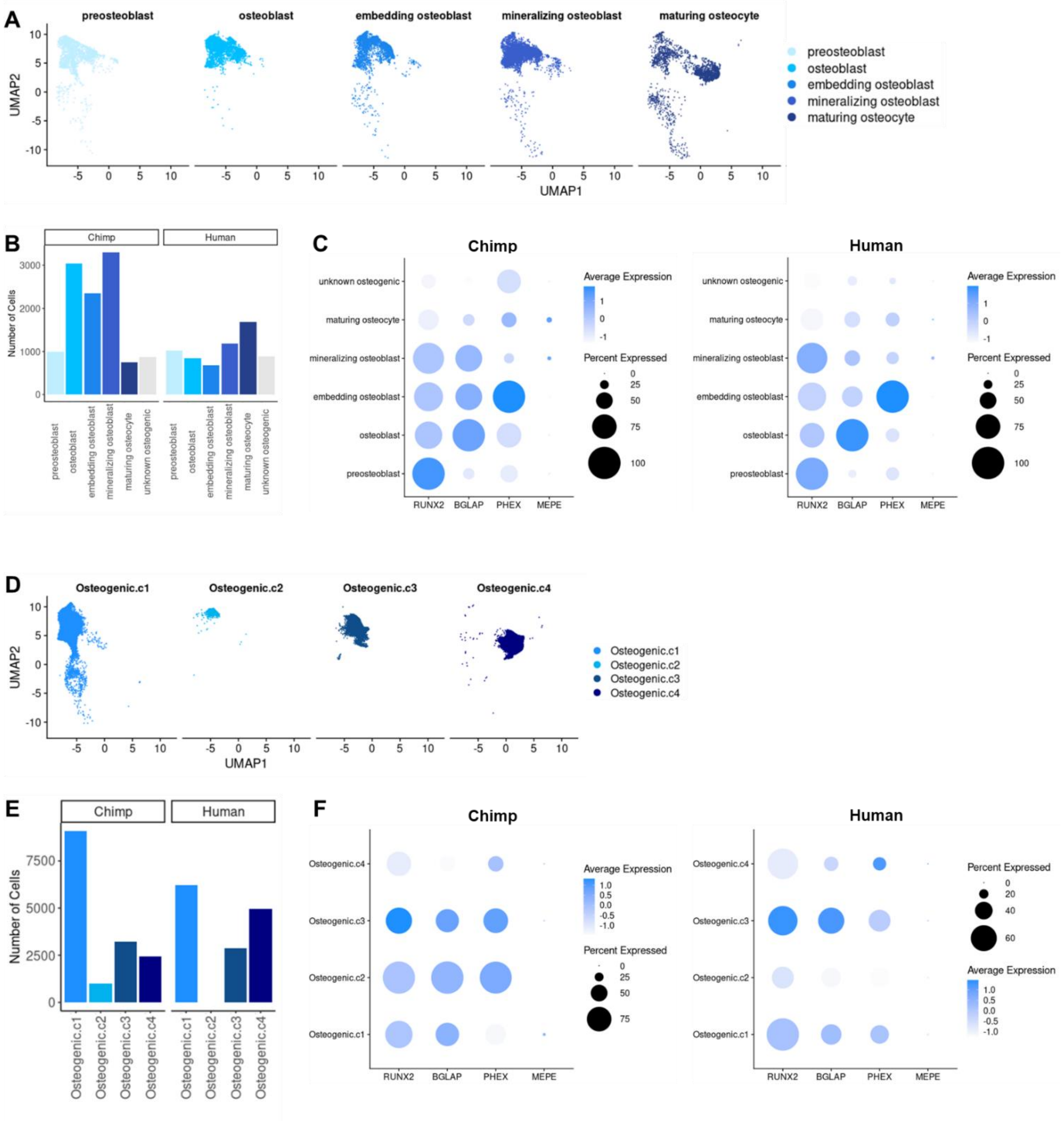

**Comparison of Osteogenic Cell Classification Schemes.** Plots compare (A-C) the osteogenic ad hoc assignment method to (D-F) the osteogenic unsupervised cluster method (resolution=0.50). (A) UMAP dimensional reduction plots of scRNA-seq data with cells labeled by the stage of osteogenesis to which they were classified via the osteogenic ad hoc assignment method. (B) Bar plot depicting the number of chimpanzee and human cells for each osteogenic ad hoc assignment. (C) Dot plots depicting the scaled average expression (dot color intensity) and the proportion of cells expressing each gene (dot size) of candidate genes (x-axis) for each osteogenic ad hoc assignment (y-axis). (C) UMAP dimensional reduction plots of scRNA-seq data with cells labeled by the osteogenic unsupervised cluster to which they were assigned. (D) Bar plot depicting the number of chimpanzee and human cells for each osteogenic cluster. (E) Dot plots depicting the scaled average expression (dot color intensity) and the proportion of cells expressing each gene (dot size) of candidate genes (x-axis) for each osteogenic cluster (y-axis).

**Figure S4.2**

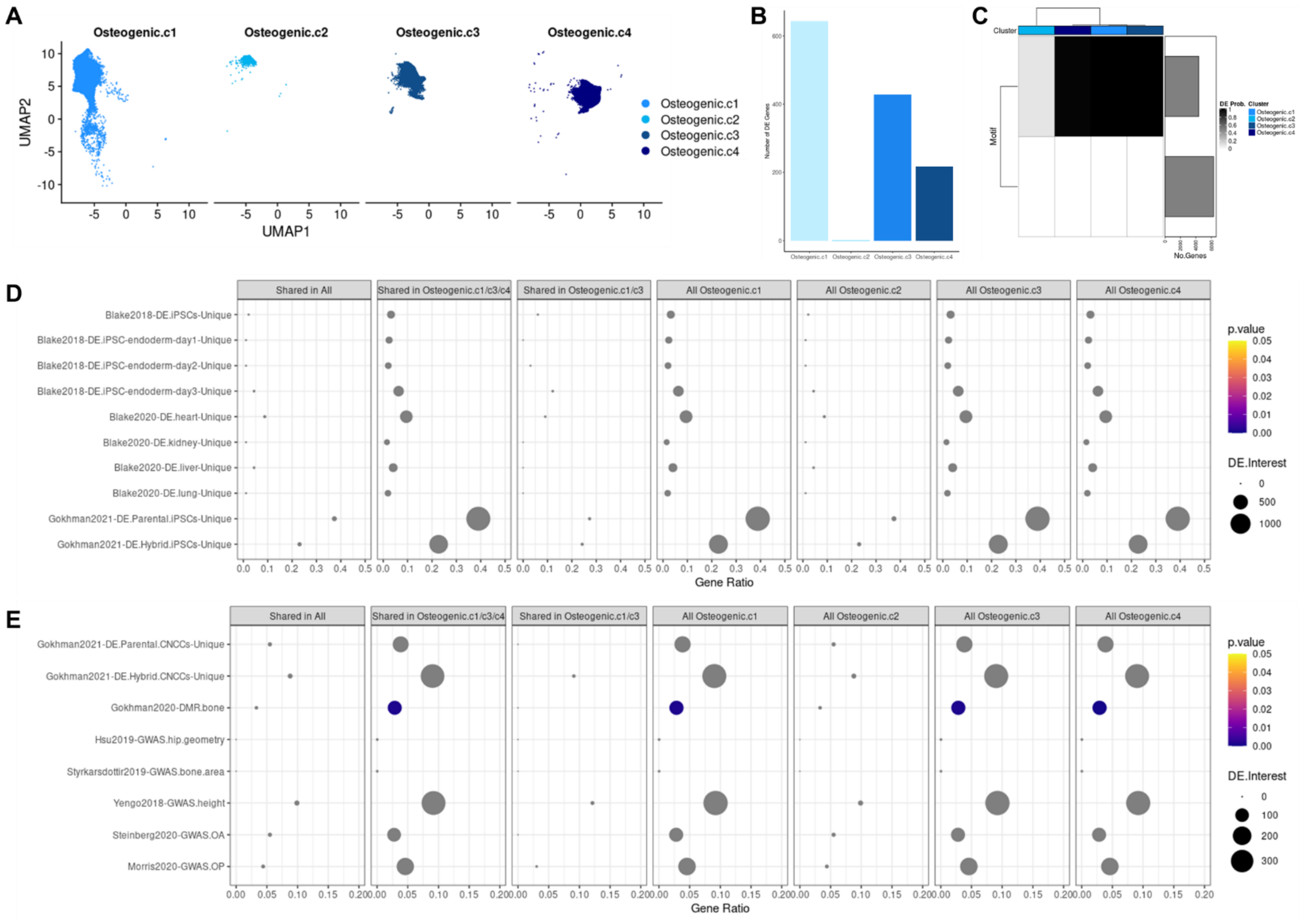

**Interspecific DE in Osteogenic Unsupervised Cluster Cell Annotations.** (A) UMAP dimensional reduction plot of scRNA-seq data with cells labeled by the osteogenic unsupervised cluster (resolution=0.50) to which they were assigned. (B) Bar plot showing the number of interspecific DE genes identified for each osteogenic cluster using standard methods. (C) Correlation motifs based on the probability of differential expression between species for each osteogenic cluster with the number of genes assigned to each motif shown in the bar plot on the right and the posterior probability that a gene is DE shown by the shading of each box. (D-E) Enrichment of external DE gene sets among Cormotif interspecific DE genes identified for each osteogenic cluster with the p-value (p.value), the number of DE genes overlapping an external gene set (DE.Interest), and the ratio of overlapping to non-overlapping DE genes for a given external gene set (GeneRatio) denoted.

**Figure S4.3**

A

| Combination of 8 Genes | | RUNX2 | BGLAP | CAPG | DMP1 | PHEX | MEPE | SOST | HYOU1 |
|---|---|---|---|---|---|---|---|---|---|
| 0 | not osteogenic | - | - | - | - | - | - | - | - |
| 1 | preosteoblast | + | - | - | - | - | - | - | - |
| 2 | osteoblast | [+/-] | + | - | - | - | - | - | - |
| 3 | embedding osteoblast | [+/-] | [+/-] | + or | + or | + & | - | - | - |
| 4 | mineralizing osteoblast | [+/-] | [+/-] | [+/-] | [+/-] | [+/-] | + | - | - |
| 5 | mature osteocyte | [+/-] | [+/-] | [+/-] | [+/-] | [+/-] | [+/-] | + or | + |

B

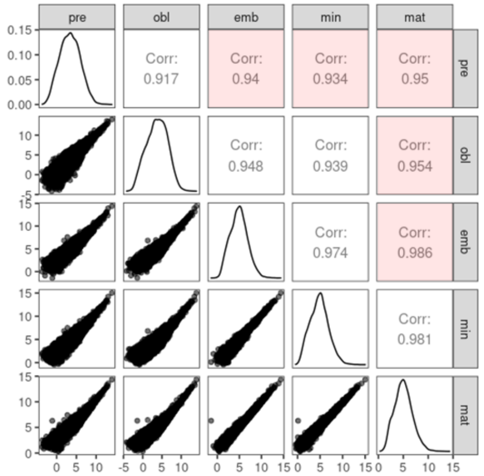

C

| Combination of 6 Genes | | RUNX2 | BGLAP | DMP1 | PHEX | MEPE | SOST |
|---|---|---|---|---|---|---|---|
| 0 | not osteogenic | - | - | - | - | - | - |
| 1 | preosteoblast | + | - | - | - | - | - |
| 2 | osteoblast | [+/-] | + | - | - | - | - |
| 3 | embedding osteoblast | [+/-] | [+/-] | + or | + & | - | - |
| 4 | mineralizing osteoblast | [+/-] | [+/-] | [+/-] | [+/-] | + | - |
| 5 | mature osteocyte | [+/-] | [+/-] | [+/-] | [+/-] | [+/-] | + |

D

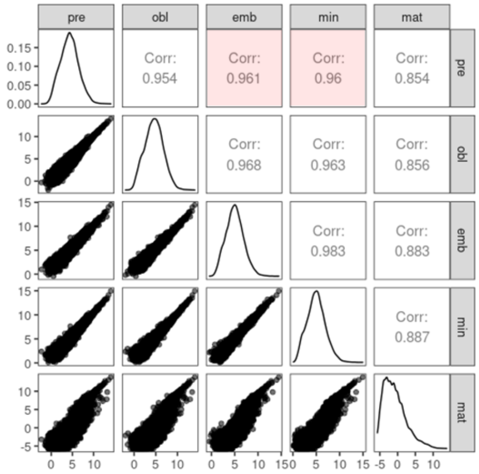

E

| Combination of 4 Genes | | RUNX2 | BGLAP | PHEX | MEPE |
|---|---|---|---|---|---|
| 0 | not osteogenic | - | - | - | - |
| 1 | preosteoblast | + | - | - | - |
| 2 | osteoblast | [+/-] | + | - | - |
| 3 | embedding osteoblast | [+/-] | [+/-] | + | - |
| 4 | mineralizing osteoblast | [+/-] | [+/-] | [+/-] | + |
| 5 | maturing osteocyte | - | - | [+/-] | + |

F

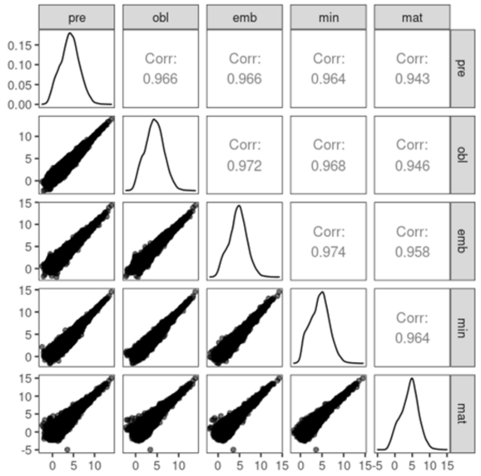

**Comparison of Osteogenic Ad Hoc Assignment Methods.** (A,C,E) Schematic for each considered osteogenic ad hoc assignment method and (B,D,F) pairwise correlations of pseudobulk counts for each gene between osteogenic ad hoc assignments. Correlation values highlighted in red indicate a deviation from the expected pattern. (A-B) Plots of the osteogenic ad hoc assignment method using 8 candidate genes. (C-D) Plots of the osteogenic ad hoc assignment method using 6 candidate genes. (E-F) Plots of the osteogenic ad hoc assignment method using 4 candidate genes.

**Figure S5.1**

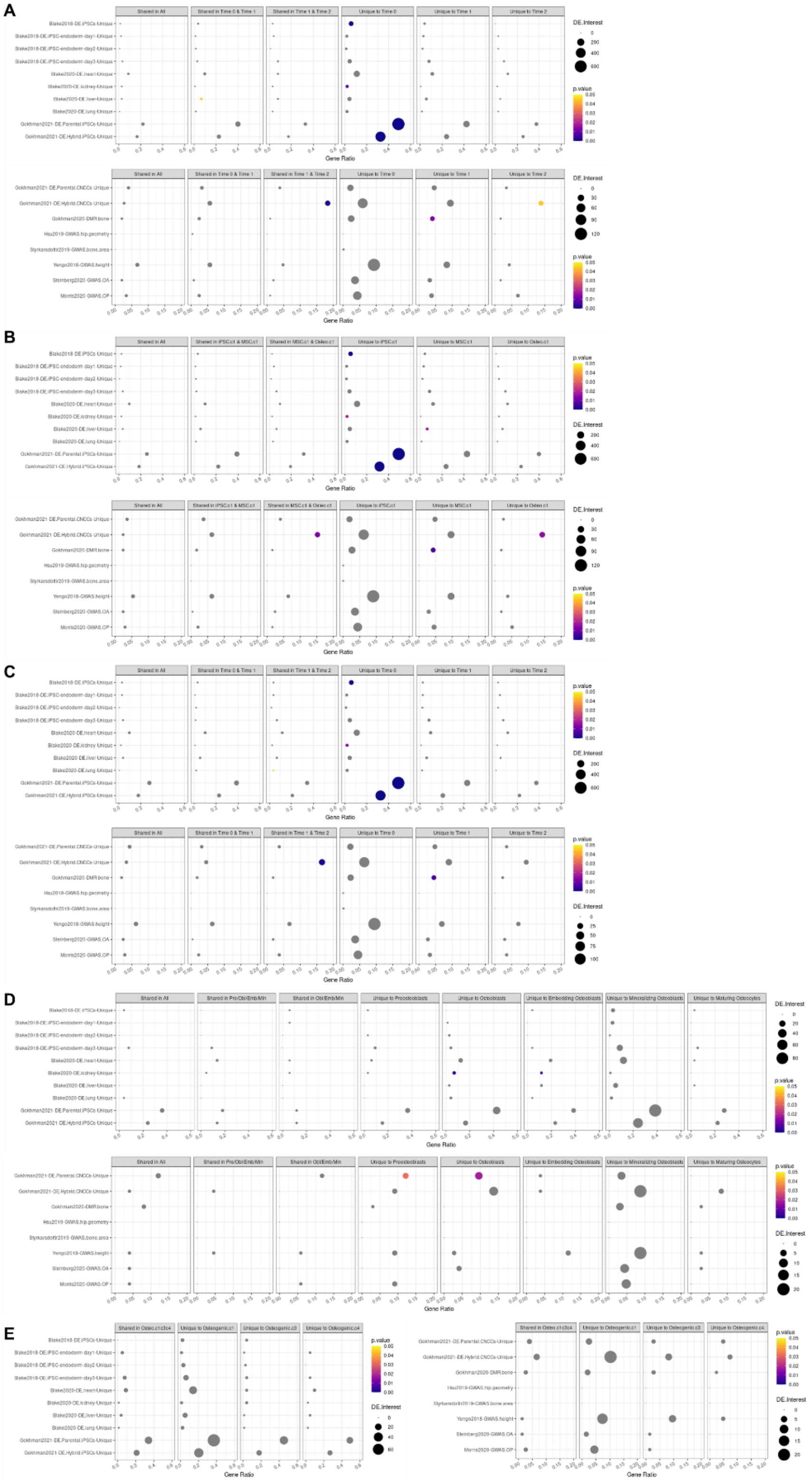

**Gene Set Enrichments in Standard Interspecific DE Genes.** Enrichment of external DE gene sets among standard interspecific DE genes (FDR<0.01) identified for given cell classifications with the p-value (p.value), the number of DE genes overlapping an external gene set (DE.Interest), and the ratio of overlapping to non-overlapping DE genes for a given external gene set (GeneRatio) denoted. (A) Enrichments across stages of differentiation. (B) Enrichments across general unsupervised clusters (resolution=0.05). (C) Enrichments across general ad hoc assignments. (D) Enrichments across osteogenic ad hoc assignments. (E) Enrichments across osteogenic unsupervised clusters (resolution=0.50).

**Figure S5.2**

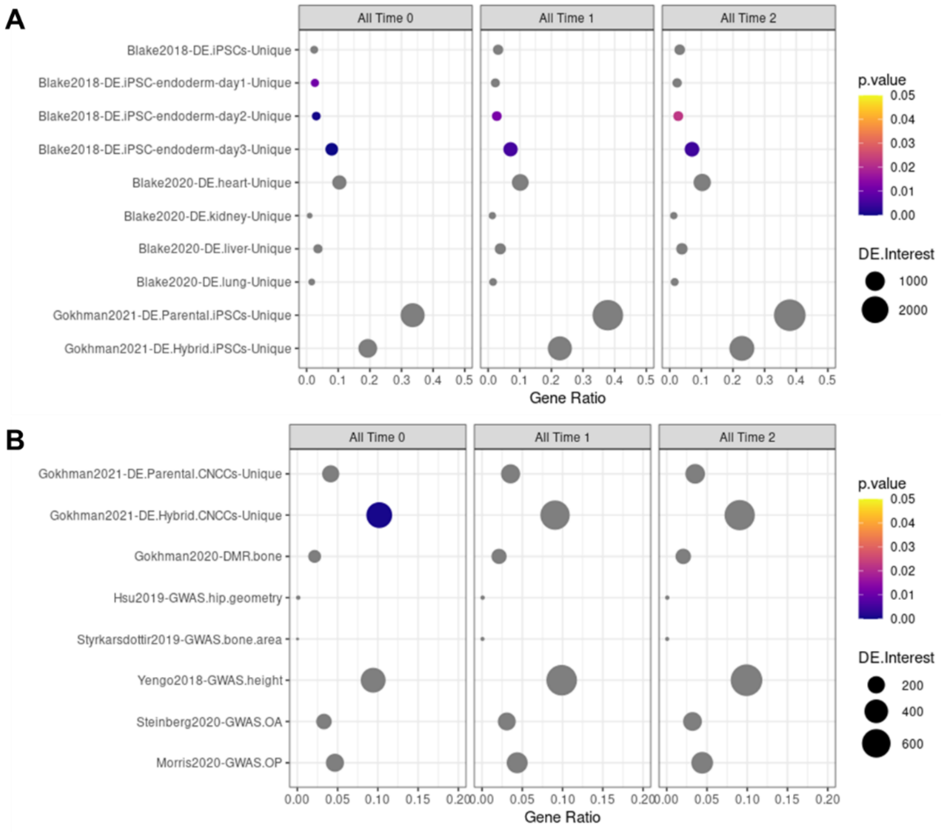

**Gene Set Enrichments in All Cormotif Interspecific DE Genes.** (A-B) Enrichment of external DE gene sets among Cormotif interspecific DE genes identified for each stage of differentiation for validation (A) and functional interpretation (B) with the p-value (p.value), the number of DE genes overlapping an external gene set (DE.Interest), and the ratio of overlapping to non-overlapping DE genes for a given external gene set (GeneRatio) denoted.

**Figure S5.3**

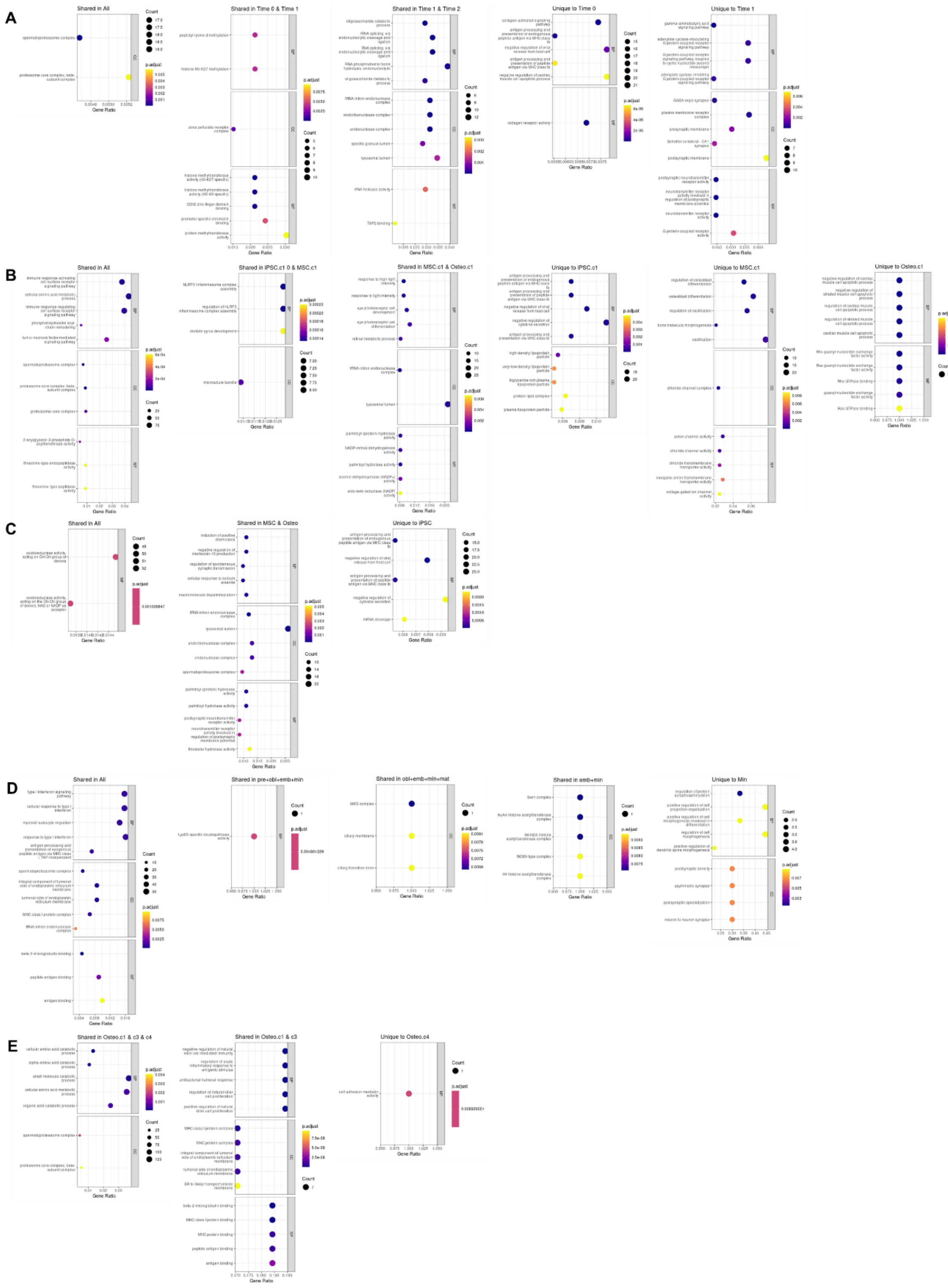

**GO Enrichments in Cormotif Interspecific DE Genes.** Enrichment of GO functional categories among Cormotif interspecific DE genes identified for a given cell classification. The top 5 GO functions identified in biological processes (BP), cell components (CC), and molecular functions (MF) are displayed along with the adjusted p-value (p-adjust), the number of marker genes overlapping a GO function (Count), and the ratio of overlapping to non-overlapping marker genes for a given GO function (GeneRatio). (A) Enrichments across stages of differentiation. (B) Enrichments across general unsupervised clusters (resolution=0.05). (C) Enrichments across general ad hoc assignments. (D) Enrichments across osteogenic ad hoc assignments. (E) Enrichments across osteogenic unsupervised clusters (resolution=0.50).

**Figure S5.4**

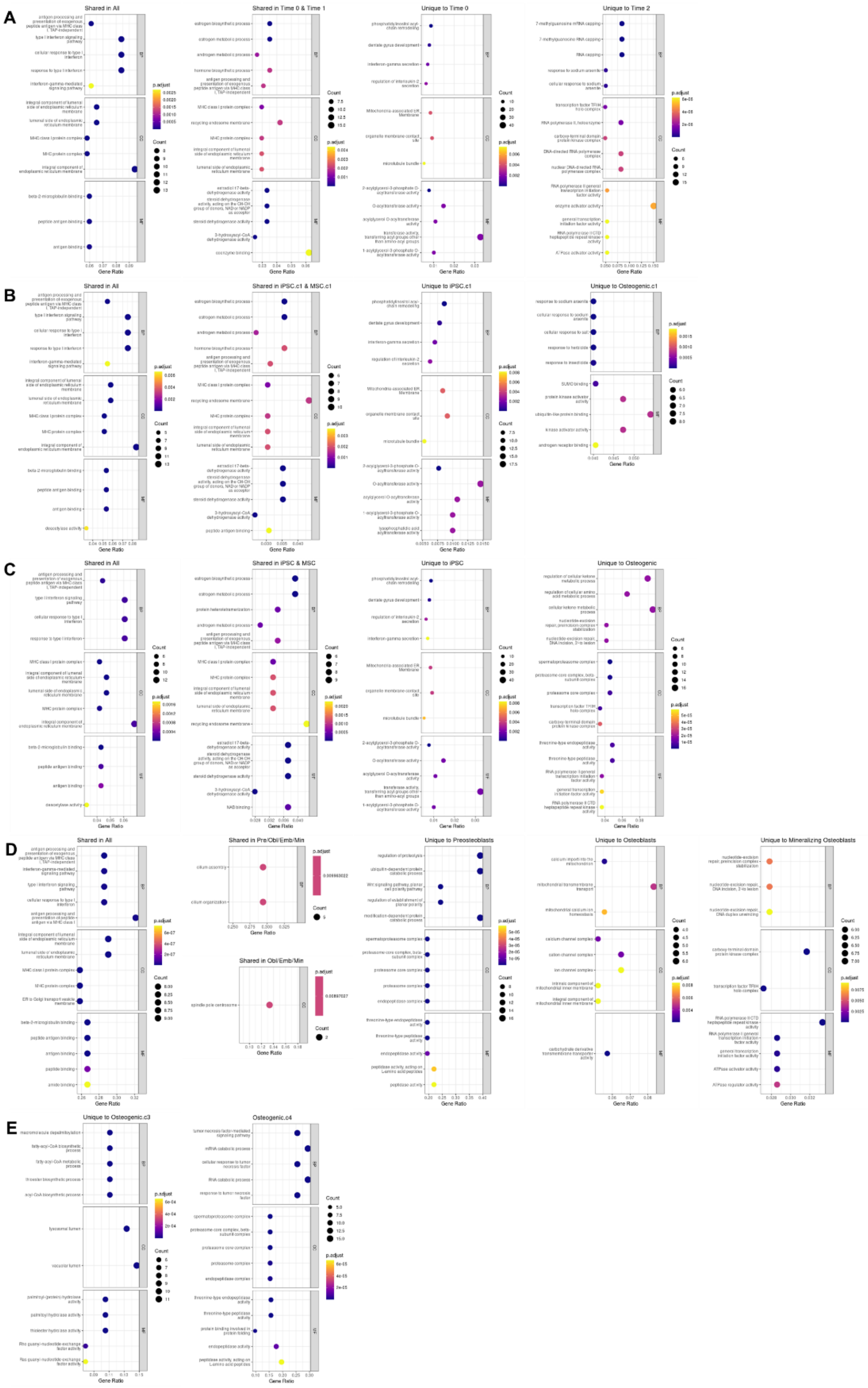

**GO Enrichments in Standard Interspecific DE Genes.** Enrichment of GO functional categories among standard interspecific DE genes (FDR<0.01) identified for a given cell classification. The top 5 GO functions identified in biological processes (BP), cell components (CC), and molecular functions (MF) are displayed along with the adjusted p-value (p-adjust), the number of marker genes overlapping a GO function (Count), and the ratio of overlapping to non-overlapping marker genes for a given GO function (GeneRatio). (A) Enrichments across stages of differentiation. (B) Enrichments across general unsupervised clusters (resolution=0.05). (C) Enrichments across general ad hoc assignments. (D) Enrichments across osteogenic ad hoc assignments. (E) Enrichments across osteogenic unsupervised clusters (resolution=0.50).
